# Molecular Classification and Comparative Taxonomics of Foveal and Peripheral Cells in Primate Retina

**DOI:** 10.1101/428110

**Authors:** Yi-Rong Peng, Karthik Shekhar, Wenjun Yan, Dustin Herrmann, Anna Sappington, Greg S. Bryman, Tavé van Zyl, Michael Tri. H. Do, Aviv Regev, Joshua R. Sanes

## Abstract

High acuity vision in primates, including humans, is mediated by a small central retinal region called the fovea. As more accessible model organisms lack a fovea, its specialized function and dysfunction in ocular diseases remain poorly understood. We used 165,000 single-cell RNA-seq profiles to generate and validate comprehensive cellular taxonomies of macaque fovea and peripheral retina. More than 80% of >65 cell types match between the two regions, but exhibit substantial differences in proportions and gene expression, some of which we relate to functional differences. Comparison of macaque retinal types with those of mice reveals that interneuron types are tightly conserved, but that projection neuron types and programs diverge, despite conserved transcription factor codes. Key macaque types are conserved in humans, allowing mapping of cell-type and region-specific expression of >190 genes associated with 6 human retinal diseases. Our work provides a framework for comparative single-cell analysis across tissue regions and species.

## INTRODUCTION

Specialized anatomic regions and neural circuitry in the central nervous system (CNS) of primates are the result of evolutionary adaptations that underlie their unique sensory and cognitive capacities. One such capacity is high visual acuity. Most primates, including humans, see objects clearly only when they look straight at them so their image falls on a small central region of the retina called the fovea, which lies within the somewhat larger macula (**Figure 1A**). This is because the fovea, which may be the only primate-specific structure in the mammalian central nervous system, mediates high acuity vision (Bringmann et al., 2018; Provis et al., 2013). Although it occupies <1% of the retinal surface, it provides ~50% of the input to primary visual cortex (Bringmann et al., 2018; Provis et al., 2013). Diseases that selectively affect this region, such as macular degeneration and diabetic macular edema, are leading causes of human blindness in the developed world (Bourne et al., 2013). These important aspects of human retinal function and dysfunction cannot be satisfactorily studied in more accessible model organisms, including mice, because they lack a fovea (Bird and Bok, 2017).

**Figure 1:**
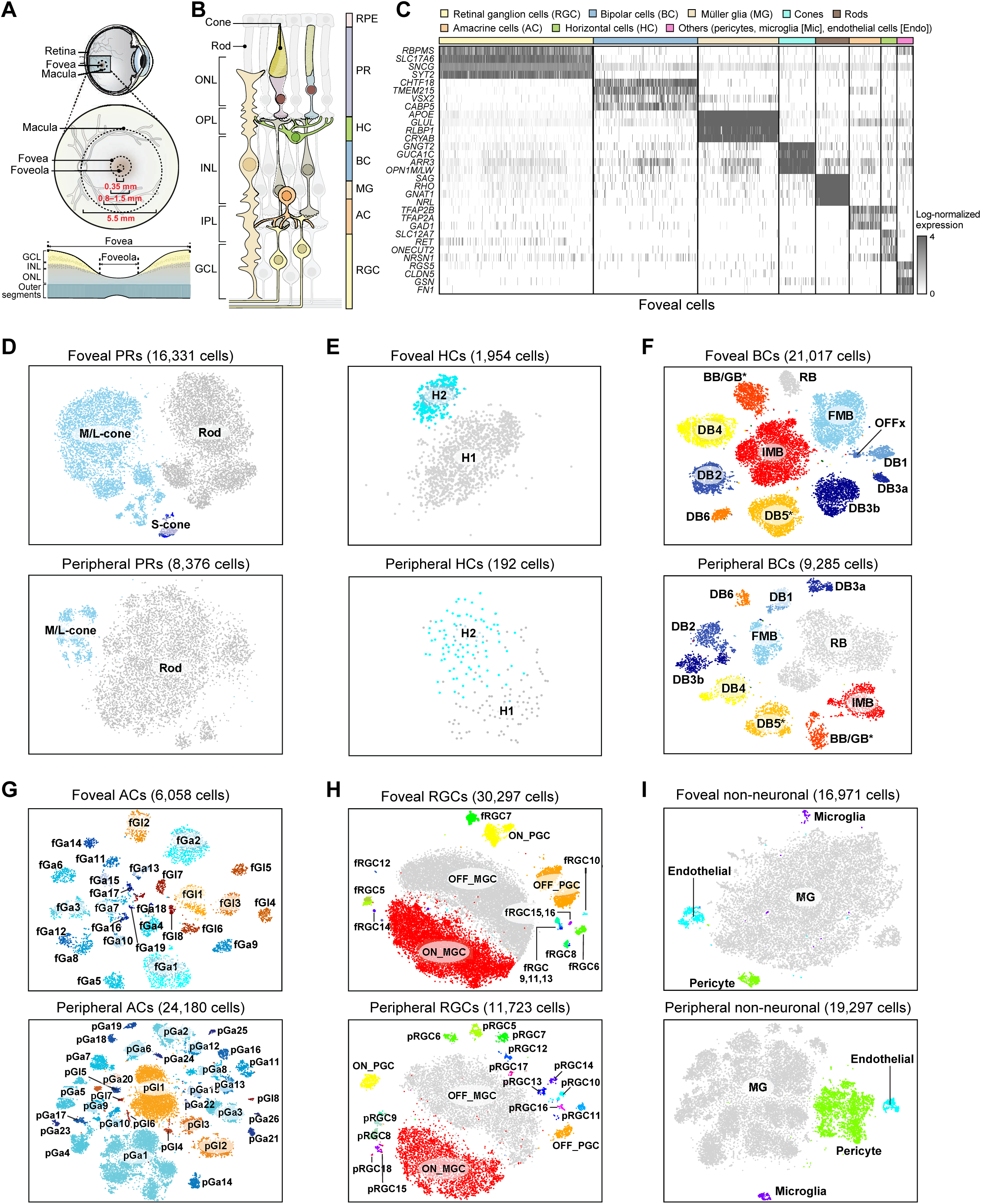
Single-cell profiling of peripheral and foveal cells from macaque retina. A. (top) Sketch of a primate eye showing position of fovea and macula. (middle) Central region at higher power indicating diameters of the foveola (the foveal pit), fovea, and macula. (bottom) Sketch of a section through macaque fovea, showing foveal pit (red arrow) and displacement of ganglion cell layer (GCL) and inner nuclear layer (INL) cells towards the foveal wall. The displacement leads to a multilayered GCL in the remainder of the fovea; in contrast the GCL in the peripheral retina contains only one or two layers of somata in peripheral retina. B. Sketch of peripheral retina showing its major cell classes – photoreceptors (PR), horizontal cells (HC), bipolar cells (BC), amacrine cells (AC), retinal ganglion cells (RGC) and Müller glia (MG), outer and the inner plexiform (synaptic) layers (OPL, IPL), outer and inner nuclear layers (ONL, INL) and ganglion cell layer (GCL). Each neuronal class is divisible into types based on morphological, physiological and molecular criteria. C. Expression patterns of class specific marker genes (rows) (**Table S1**) in single foveal cells (columns). Cells are grouped by their class (color bar, top). Plot shows randomly selected 10% of total cells. These signatures were used to separate peripheral cells into classes (not shown). D-I. t-distributed stochastic neighbor embedding (tSNE) visualization of foveal (top panel) and peripheral (bottom panel) PRs (D), HCs (E), BCs (F), ACs (G), RGCs (H), non-neuronal cells (I). Individual cells (points) are colored by their cluster assignments. Cluster labels, corresponding to *post hoc* assigned types, are indicated. Although substructure is visible in D and I, this reflected batch effects and we were unable to detect subtypes by reanalysis. Foveal (f) and peripheral (p) AC clusters are divided into GABAergic (Ga) and Glycinergic (Gl) groups, and then numbered from largest to smallest cell number within each group. For RGCs, OFF and ON midget (MGC) and parasol (PGC) ganglion cells are labeled; clusters 5+ numbered from largest to smallest in the fovea (f) and periphery (p), respectively.

Basic aspects of retinal structure and function are shared by all vertebrate species (**Figure 1B**), and between the fovea and the peripheral retina. Photoreceptors (PR, rods and cones) convert light into electrical signals, which are transferred to interneurons (horizontal, bipolar and amacrine cells; HCs, BCs and ACs). Interneurons process the information and deliver it to retinal ganglion cells (RGCs), which send axons through the optic nerve to the rest of the brain (Masland, 2012). Selective connectivity among multiple AC, BC and RGC types, along with type-specific intrinsic differences, endow each RGC type with selective responsiveness to specific visual features (Gollisch and Meister, 2010; Sanes and Zipursky, 2010).

Despite a conserved fundamental plan, there are numerous differences between the fovea and the peripheral retina (Bringmann et al., 2018; Greeff, 1874; Sinha et al., 2017). For example, many RGCs in the fovea are supplied by a single PR, which maximizes acuity, whereas peripheral RGCs are supplied by dozens to hundreds of PRs, enhancing sensitivity (**Figure S1A**). In addition, the fovea is rich in cones, which mediate color and contrast-sensitive vision in well-lit conditions, whereas peripheral retina is dominated by rods, which can function in dim light levels, further enhancing sensitivity (night vision). Beyond compositional differences, cell types in the two regions exhibit distinct morphologies and physiological functions (**Figure S1A**) (Greeff, 1874; Kolb and Marshak, 2003; Sinha et al., 2017). Because molecular information is lacking for most primate retinal cell types, the underpinnings of these differences are little explored. For example, it is unclear the extent to which differences stem from cell types unique to one region, from altered proportions across the same set of types, or from region-specific molecular programs within shared types.

Lack of molecular knowledge has also precluded systematic assessment of the conservation of retinal cell types between primates and model organisms for which greater molecular and functional information is available. As evolutionary distances grow, identification of orthologous cell types becomes challenging in the absence of such knowledge. The retina provides an opportunity to assess molecular strategies for establishing orthologies. In mouse retina, there are >110 neuronal types, with molecular signatures as well as morphological and physiological characterization available for many of them (Sanes and Masland, 2015; Shekhar et al., 2016). In contrast, although many primate retinal types have been identified (~60 to date), they have been defined largely by morphological criteria e.g. (Masri et al., 2016; Morgan and Wong, 2008; Santina and Ou, 2018), which has not proven to be a satisfactory means of matching them to mouse types. Molecular characterization could provide a lingua franca to enable mapping, which would not only inform how common model organisms relate to primates, including humans, but also aid in understanding the evolution of cell types and their associated expression programs.

To systematically address these questions, we generated and analyzed an atlas of >165,000 single-cell RNA-seq (scRNA-seq) profiles from the fovea and peripheral retina of adult crab-eating macaques (*Macaca fascicularis*), a widely used primate model in vision studies (**Figure S1B**). We identified and molecularly characterized >65 cell types in each region, providing novel molecular markers for most types, and validated many by pairing them with cell morphology. We used a comparative analytic framework to match cell types between the fovea and the periphery, showing that the two regions share a common repertoire of types, but exhibit marked differences in cell type proportions and cell-intrinsic expression programs, which likely mediate their functional differences. We then assessed the conservation of macaque retinal cell types with those in other species. We found a tight correspondence between mouse and macaque PRs, BCs and ACs, but a divergence in RGC types, despite a conserved transcription factor code. Key macaque types and molecular features are also conserved in marmosets and humans. Finally, we used our atlas to determine the cell- and region-specific expression of >190 genes implicated in 7 complex diseases that cause human blindness, demonstrating striking patterns of region- and cell type-specific expression in macaques, which are, however, not conserved in mice. Our study provides a general framework for using molecularly derived taxonomies of cell types to understand regional and species specializations in the nervous system and other organs.

## RESULTS

### Comprehensive molecular taxonomy of the primate retina using scRNA-seq

We generated cell atlases of the foveal and the peripheral regions of the primate retina using droplet-based scRNA-seq (Zheng et al., 2017) (see **Methods**), yielding high-quality cell profiles from 92,628 foveal, and 73,053 peripheral retinal cells. For foveal samples, cells were dissociated from 0.5-1.5 mm diameter samples obtained from 4 adult macaques (1 fovea per animal), and profiled without further processing. Peripheral retinal cells were obtained from 4 adults, one of which had also provided a foveal sample. Because the peripheral retina is dominated by rod PRs (~80% of total cells), we used magnetic columns to deplete rods (CD73+) or enrich RGCs (CD90+) (**Figure S1B**, see **Methods**). In parallel, we assembled a retina-specific transcriptome, which substantially improved the mapping of scRNA-seq reads compared to existing references (**Figures S1C-E**), emphasizing the importance of high-quality tissue-specific transcriptome for effective scRNA-seq analyses.

To maximize our ability to distinguish cell types consistently across biological replicates, we adopted an informatics approach in which we first grouped cells into the 6 major retinal classes (RGCs, BCs, PRs, ACs, HCs, and non-neuronal cells) across all samples based on known class-specific gene signatures (**Figures 1C, S1F, see Table S1**), and then iteratively clustered cells within each class separately for the fovea and the periphery, building on our earlier methods (Shekhar et al., 2016).

Altogether, we distinguished 64 foveal (3 PR, 2HC, 12 BC, 27 AC, 16 RGC, 4 non-neuronal), and 71 peripheral (2 PR, 2 HC, 11 BC, 34 AC, 18 RGC and 4 non-neuronal) clusters (**Figures 1D-I**); all clusters were found in each animal (**Figures S1G-I**). Non-neuronal cells were comprised of Müller Glia, pericytes, endothelial cells and microglia (**Figure 1I**). There are no oligodendrocytes in normal retina, and astrocytes, which predominantly ensheath RGC axons in the nerve fiber layer (Vecino et al., 2016), were not recovered in our samples. Using differential expression analysis, we identified molecular markers for each cluster within the foveal and peripheral samples. We then used this information to assign clusters to individual retinal cell types. We first describe the types, then return to document differences between corresponding foveal and peripheral types.

### L and M cones are distinguished only by expression of opsin paralogs

We first analyzed PRs. Retinas of trichromatic primates, including macaques and humans, contain rods plus three cone types that preferentially detect long (L, red), medium (M, green) and short (S, blue) wavelengths, based on their expression of L, M and S opsins (*OPN1LW, OPN1MW, OPN1SW*) respectively. S cones were readily distinguished, whereas M and L cones mapped to a single cluster (**Figure 1D**). *OPN1MW* and *OPN1LW* have ~98% identical coding sequences (Onishi et al., 2002) and are not distinguished in the macaque reference genome. However, by examining relevant *OPN1LW/MW* reads for diagnostic single-nucleotide polymorphisms, we could assign ~70% of M/L cones as either L or M (**Figure S2A-C,E**).

Remarkably, no genes other than *OPN1MW* or *OPN1LW* were differentially expressed (DE) between the M and L cones (**Figures 2A, S2D**), supporting a model in which stochastic choice between the neighboring *OPN1MW* and *OPN1LW* genes assigns otherwise identical cones to one of two groups (Wang et al., 1999); it is less consistent with models that imply distinct transciptional programs in the two cone types (Lee et al., 2012). In contrast, many genes distinguished M/L from S cones and rods, and S cones from rods, several of which we validated using fluorescent *in situ* hybridization (FISH, **Figures 2B,C and S2F**).

**Figure 2:**
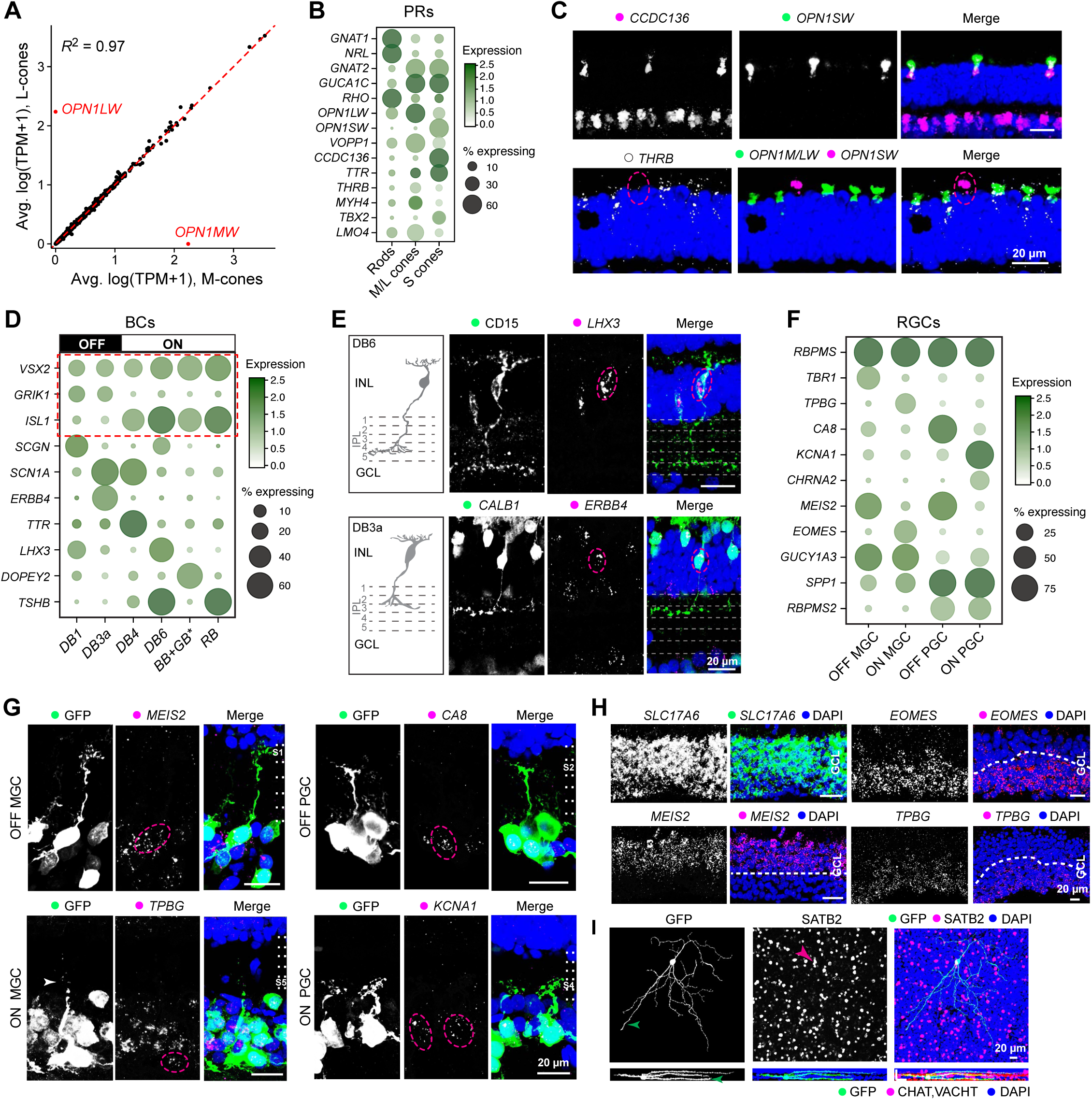
Matching scRNA-seq clusters to neuronal types of the primate retina. A. Comparison of average transcriptional profiles of foveal M-cones and L-cones. Each dot corresponds to a gene. No genes other than *OPN1MW* and *OPN1LW* differ significantly in expression levels (>1.2 fold at p<0.01, MAST test) between the two cone types. B. Dot plot showing expression of genes (rows) that distinguish photoreceptor (PR) types (columns), common to both the fovea and the periphery. The size of each circle is proportional to the percentage of cells expressing the marker (≥1 UMI), and its intensity depicts the average transcript count within expressing cells. C. Validation of S cone specific gene *CCDC136* (upper) and M/L cone specific gene *THRB* (lower) by double and triple FISH with *OPN1SW* (S-opsin) and *OPN1MW/LW* (M/L-opsin). Circle highlights an S cone. D. Gene expression patterns of type-enriched markers for selected BC types. Red box highlights a pan-BC, a pan-ON-BC and a pan-OFF-BC marker. Representation as in panel B. See **Figure S2G** and **Table S2** for lists. E. Validation of new markers for two BC types by FISH combined with immunostaining for cell morphology. (top) DB6 (circle) cells, known to be CD15-positive, also express *LHX3*. (bottom) DB3a cell (circle) is CALB1*+ERBB4+*. IPL sublaminae (S1-S5) are demarcated by dashed lines. Sketches redrawn from (Tsukamoto and Omi, 2015, 2016). F. Expression patterns of genes selectively enriched among foveal ON and OFF MGCs and PGCs. Representation as in panel B. G. Validation of markers for MGCs and PGCs from panel F, combining FISH with viral labeling (GFP) to show RGC morphology in the fovea. H. Somata of ON MGCs (*TPBG+* and *EOMES+*) and OFF MGCs/PGCs (*MEIS2+*) are localized to the inner and outer halves (divided by dashed lines) of the ganglion cell layer, respectively, proximal to the fovea. **I**. SATB2 positive SBC labeled by GFP-expressing virus. Arrowheads indicate the axon (green) and soma. Bottom panels are rotations to show bistratified dendritic lamination. Scale bar is 20 μm. DAPI staining is blue in C, E, G-I.

### Molecular classification and validation of interneuronal types

Using a combination of prior knowledge and new histological analyses, we next associated molecularly defined clusters of foveal and peripheral interneurons (BCs, ACs, and HCs) with individual cell types.

BCs, the major excitatory interneurons of the retina, are divided into those that receive predominant input from cones or rods, and cone BCs are further divided into those excited (ON BCs) or inhibited (OFF BCs) by light; all rod BCs are ON (Euler et al., 2014). Twelve BC types (1 rod, 5 OFF cone and 6 ON cone) have been previously reported in macaque peripheral retina based on morphological and immunohistochemical analyses (Joo et al., 2011; Tsukamoto and Omi, 2015, 2016).

We identified 12 foveal BC and 11 peripheral BC clusters; 10 mapped 1:1 to known types (**Table S2**), and supervised methods separated another cluster into two known types in both regions (**Figure S2H**). One foveal cluster corresponded to a type not described previously. Based on gene expression, it is likely an OFF type; we provisionally call it OFFx (**Figure S2I**). We identified markers for each BC type and validated several of them (**Figures 2D,E, S2G,J-O**).

HCs and ACs are the main source of inhibitory input to retinal circuitry. We identified 2 clusters of HCs in both fovea and periphery (**Figure 1E**), and confirmed that the corresponding cluster-specific markers labeled distinct HC subsets in tissue. We also validated *SPP1*, which encodes for the secreted protein osteopontin, as a pan-HC marker (**Figure S5D**). We similarly identified 27 foveal AC clusters, and 34 peripheral AC clusters, including both GABAergic and glycinergic subfamilies (**Figures 1G, S4A, S5E, Table S2**). ACs remain the least characterized retinal class across all vertebrate species. However, we tentatively assigned six clusters to known mouse types based on their expression of orthologous markers, including Starburst, VIP+, SEG, catecholaminergic, glutamatergic (VGluT3+), and SEG ACs (**Figures S4A and S5E**). In addition, we found the expression of neuropeptides in subsets of GABAergic AC types, which is consistent with previous studies (**Table S2**).

### Markers for midget and parasol RGCs

Morphological studies have distinguished 17-20 RGC types in primate retina (Dacey, 2004; Masri et al., 2016). ON and OFF Midget RGCs (MGCs), the smallest and most abundant, comprise ≥85% of foveal RGCs; larger ON and OFF parasol RGCs (PGCs) comprise ~10%; small bistratified RGCs (SBCs) and ~12 types with larger dendritic arbors account for the remaining ~5%. MGCs and PGCs are also the most abundant RGC types in the periphery, together accounting for ~70% of peripheral RGCs (Dacey, 2004). Axons from MGCs, PGCs and SBCs innervate different layers of the lateral geniculate nucleus and, indirectly, different sets of cortical targets (Dacey, 2004).

We identified 18 peripheral and 16 foveal RGC clusters (**Figure 1H**). Based on their abundance, we tentatively identified one pair of clusters as ON and OFF MGCs (~85% of RGCs in fovea, 83% in periphery) and a second pair as ON and OFF PGCs (11% in fovea, 4.8% in periphery). We identified markers for these types, including *TBR1* (OFF MGC), *TPBG* (ON MGC), *CHRNA2* (ON PGC) and CA8 (OFF PGC). Among MGCs and *PGCs, SPP1* and *RBPMS2* are expressed by both PGC types and *GUCY1A3* by both MGC types (**Figure 2F**, **S3C-O**). We confirmed the identities of these four RGC types by combining *in situ* hybridization for selective markers with viral labeling to visualize morphology (**Figures 2G, S3B,P,Q**). Interestingly, somata expressing transcription factors selective for the major ON and OFF RGC types (MGCs and PGCs) were segregated into outer and inner halves of GCL, suggesting a molecular basis for the laminar organization of physiologically distinct subclasses (Perry and Silveira, 1988) (**Figures 2H and S3P**). We were unable to further partition MGCs and PGCs by supervised methods.

Three peripheral RGC clusters expressed *OPN4* (melanopsin), a marker of intrinsically photosensitive RGCs (ipRGCs) (Do and Yau, 2010). We identified markers for the remaining clusters, including *SATB2* as a marker for SBCs and two other types (**Figures 2I and S3A**). To our knowledge, these are the first molecular markers for the major RGC types of the primate retina (**Figure S3A**).

### Most retinal cell types match 1:1 between the fovea and the periphery, but differ in proportion

We devised a computational approach to relate foveal and peripheral clusters based on their molecular expression patterns. While it was straightforward to match PR types (rods and cones) and non-neuronal types (Müller glia, pericytes, endothelial cells and microglia) across regions based on known markers, HCs, BCs, ACs and RGCs required a more systematic approach. We used a multi-class learning framework that associated each foveal cell with a corresponding peripheral identity, and then examined the extent to which clusters across the two regions mapped specifically (see **Methods**).

Remarkably, 77% of foveal clusters exhibited a 1:1 mapping with peripheral clusters including 10 of 12 BC, 2 of 2 HC, 20 of 27 AC and 14 of 16 RGC clusters (**Figures 3A-D**). The matches between the foveal and the peripheral clusters was significantly higher than predicted by chance, as quantified by the Adjusted Rand Index (ARI) values, a measure of similarity between data clusterings. Six foveal clusters (RGC clusters fRGC11 and fRGC14, and AC clusters fGa3, fGa7, fGa13 and fGl8) could be partitioned because they each mapped to two closely related peripheral clusters. Notably, there were only three instances of multiple (≤ 3) foveal clusters mapping to a single peripheral cluster. Most instances of multi-mapping were for AC clusters, which likely reflects a biased frequency distribution in the peripheral samples, which were not optimized for AC recovery (**Figures S4B**).

**Figure 3:**
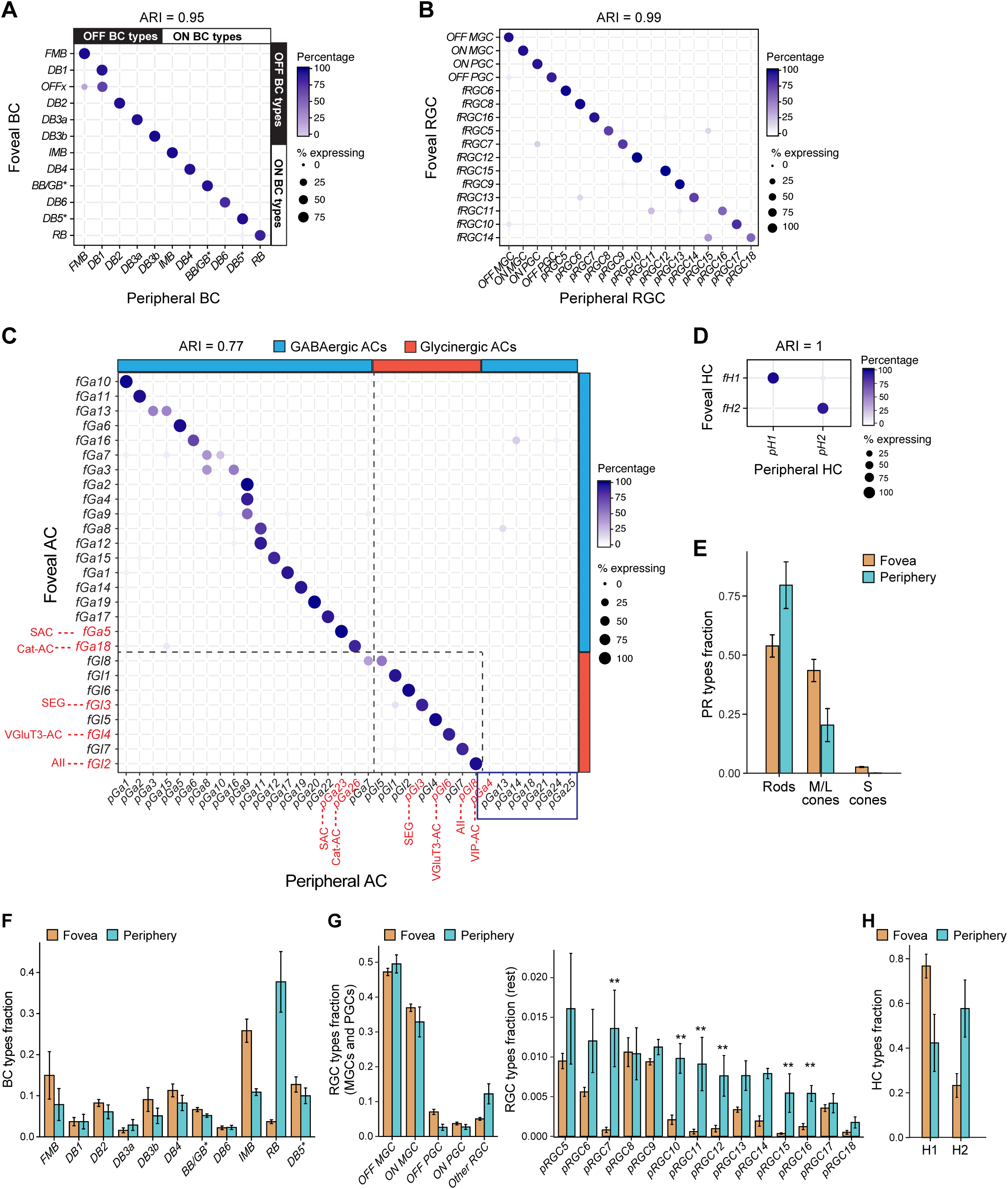
Correspondence between foveal and peripheral clusters. A-D. Transcriptional correspondence between foveal and peripheral clusters, summarized as “confusion matrices.” Circles and colors indicate the percentage of cells of a given foveal cluster (row) assigned to a corresponding peripheral cluster (column) by the classification algorithm trained on peripheral cells. A, BCs; B, RGCs; C, ACs; D, HCs. In panel A, bars on the top and right border mark ON and OFF BC subgroups. In C, bars mark GABAergic and Glycinergic AC subsets; key known types are labeled in red and types that might be periphery-specific are highlighted by blue box in C. **Figure S4A** provides molecular markers for AC clusters. In panels A-D, the extent of 1:1 cluster matches are quantified by values of the Adjusted Rand Index (ARI), which range from close to 0 (random) to 1 (perfect 1:1 match). Empirical ARI values were highly significant for all classes, as compared to null ARI values (mean ± SD) from random associations, which were as follows. BCs: 10^−5^ ± 6 × 10^−4^, RGCs: −2 × 10^−5^ ± 6 × 10^−4^, ACs: −8.4 × 10^−6^ ± 3 × 10^−4^ and HCs: 3 × 10^−6^ ± 2 × 10^−4^. E-H. Comparison of cell type proportions between the fovea and the periphery (mean±SD). Error bars represent standard deviations computed across biological replicates. E, PRs; F, BCs; G, RGCs; H, HCs. Foveal type OFFx and DB1 are grouped together as “DB1” due to their transcriptional similarity. To facilitate direct comparison of RGCs, each foveal type is assigned a peripheral identity from B. For ACs, see Figure S4A. Supervised analysis split fRGC11 and fRGC14 into two types each. RGC types underrepresented in the fovea are marked ** in G.

Quantifying the compositional similarity between the fovea and the periphery for each neuronal class suggested that differences decrease from outer to inner retina. Specifically, values of the Jensen-Shannon divergence (JSD) - a distance measure in frequency space ranging from 0 (identical cell type composition) to 1 (non-overlapping, region-specific cell types) were: PRs (0.21), BCs (0.16), ACs (0.06), and RGCs (0.035) (see **Methods**), suggesting that the major compositional differences between the fovea and periphery is within the sensors (PRs) and the earliest processors (BCs). There were numerous differences in relative proportions for 1:1 matched types (**Figures 3E-H** and **S4B, Table S2**), some consistent with previous reports, such as the predominance of rod BCs in peripheral retina, and the enrichment of cone midget bipolars (IMB and FMB) in fovea. A few types appeared to be highly restricted to either fovea or periphery. Six RGC types were greatly underrepresented (>4-fold) in foveal compared to peripheral samples (**Figure 3G**). In addition, 6 GABAergic AC types appeared to be periphery-specific (**Figure 3C**) and the OFFx BC type appeared to be fovea-specific (**Figure S2I**). Taken together, these results suggest that foveal and peripheral neuronal circuitry draw upon a similar but not identical “parts list” of cell types.

### Programs for phototransduction and GABAergic neurotransmission differ between matching types in the fovea and periphery

We next asked whether changes in gene expression between corresponding cell types further distinguished the fovea and periphery (**Figures 4, S4**). Indeed, we found a median of 17 ± 8 genes significantly DE between corresponding types (**Table S3**), with more differences for RGCs and non-neuronal cells (median 130 genes per type) compared to interneurons (HCs, BCs and ACs; median 13 genes per type) (**Figure 4A**).

**Figure 4:**
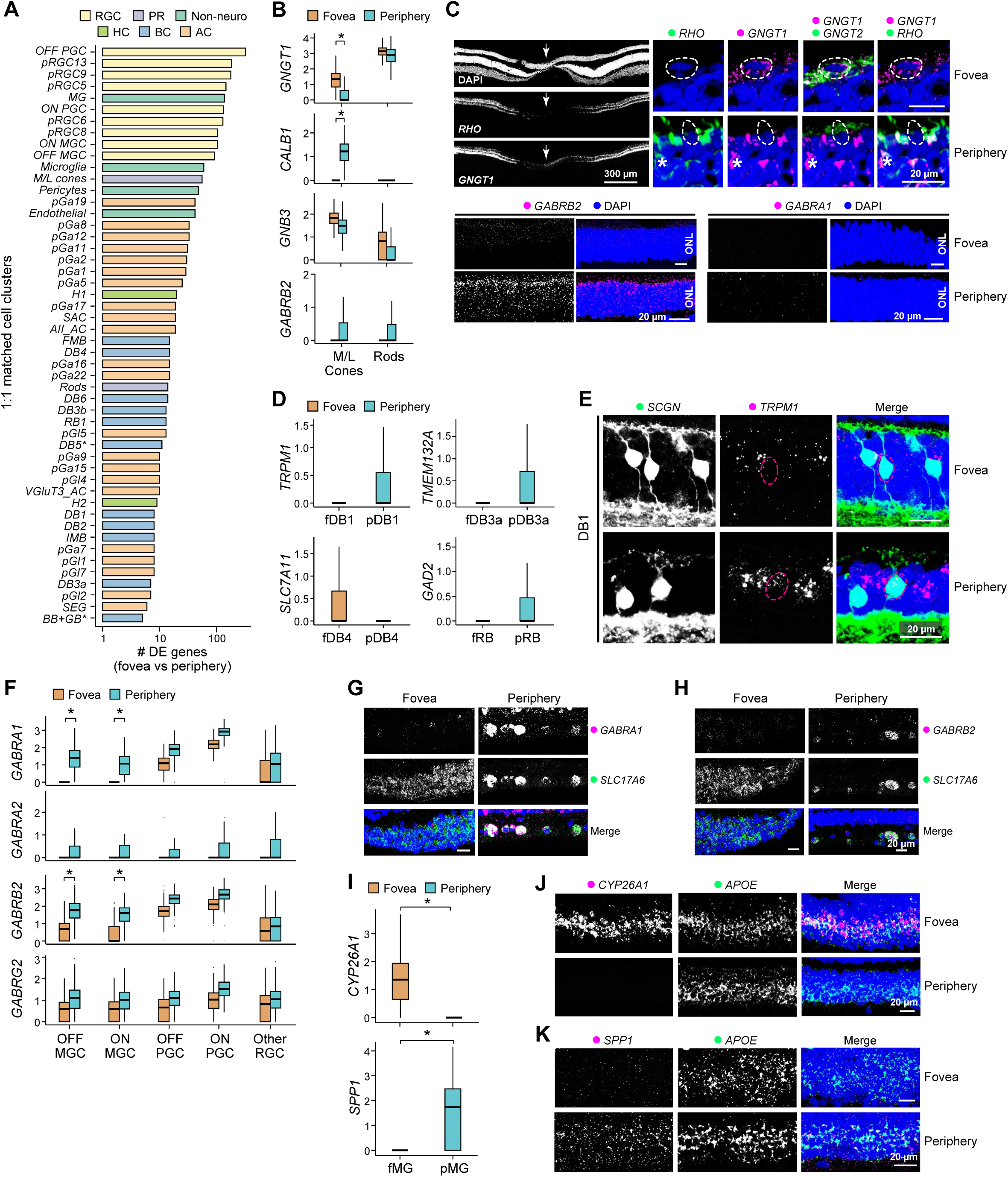
Differences in gene expression between foveal and peripheral cell types. A. Barplot showing the number of DE genes per matched cluster between the fovea and the periphery (log-fold change > 2, p < 10^−5^, MAST test). Bars are labeled based on the corresponding peripheral cluster (except known types), colored by cell class and arranged in decreasing order. Only 50 clusters that have at least 50 cells in both the fovea and periphery are shown. B. Box and whisker plots show examples of gene expression differences foveal and peripheral PRs. *p*-values were calculated using the MAST package. Black horizontal line, median; Bars, interquartile range; vertical lines, minimum and maximum. All differences between fovea and periphery in B,D,F,J are statistically significant with p<1e-19. Stars indicate >2-fold changes based on means. C. In situ validation for B. (upper) Foveal but not peripheral cones express *GNGT1*. Arrow indicates foveal center. Circles highlight cones; asterisks highlight rods (*RHO*+). (lower) Peripheral PRs express higher *GABRB2* than foveal PRs; neither type expresses *GABRA1*. D. Same as B, for BCs. E. In situ validation for D. *TRPM1* is expressed in peripheral but not foveal DB1. Both are SCGN+. F. Same as B, for RGCs. G., H. In situ validation for F. *GABRA1* (G) and *GABRB2* (H) expression in foveal and peripheral RGCs (*SLC17A6+*). I. Same as B, for MGs. J., K. In situ validation for I. *CYP26A1* (J) and *SPP1* (K) are selectively expressed by foveal and peripheral MGs respectively. *APOE* is a pan-MG marker. Scale bars, 20 and 300μm as indicated. DAPI staining is blue in C,E,G,J.

One noteworthy difference, which might underlie a functional foveal specialization, is related to phototransduction. The phototransduction component *GNGT1*, which is selectively expressed by rod PRs in mice and humans, and at higher levels by rods than cones in macaques (**Figure 4B**), is expressed at higher levels in foveal than peripheral cones, as verified by FISH and immunohistochemistry (**Figure 4C** and see below). A recent study showed that light responses are slower in macaque foveal than peripheral cones (Sinha et al., 2017), and homozygous deletion of *Gngt1* accelerates rod responses in mice (Kolesnikov et al., 2011). It is thus tempting to speculate that *GNGT1* contributes to the kinetic differences in macaques. Conversely, another phototransduction component *GNB3* is thought to be selectively expressed by cones but is also expressed at substantial levels by foveal rods (**Figure 4B**). Likewise, although the molecular profiles of foveal and peripheral BC types were extremely similar, the cation channel gene *TRPM1* is expressed in peripheral but not foveal OFF DB1 cells (**Figures 4D,E**).

Another prominent difference is that several genes involved in GABAergic neurotransmission are expressed at higher levels in peripheral types than in their foveal counterparts. They include the GABA_A_ receptor β2 subunit *GABRB2* in peripheral PRs (**Figures 4B,C**), the GABA synthetic enzyme *GAD2* in peripheral rod BCs (**Figure 4D**) (Lassova et al., 2010) and several GABA receptor subunits in multiple peripheral GC types, including MGCs and PGCs (**Figure 4F-H and S4C**). Thus, the paucity of inhibitory drive to foveal midgets observed physiologically (Sinha et al., 2017) reflects both the absence of several GABAergic AC types from fovea (**Figure 3C**) and decreased expression of GABAergic components in shared cell types. These differences suggest distinct signaling processes in the two regions.

Interestingly we also observed >100 DE genes between foveal and peripheral MGs, including *SPP1* enriched in peripheral MGs and *CYP26A1* in foveal MGs (**Figures 4I-K, S4D**). These differences provide potential substrates for differences between their interactions with neurons in the two regions (Bringmann et al., 2018; Greeff, 1874; Sinha et al., 2017).

### Tight molecular correspondence between macaque and mouse BC and AC types

While the conservation of genes across species has been extensively studied, we know less about the extent to which cell types are conserved, and such comparisons have often been challenging given the potentially rapid evolution of defining markers (Shay et al., 2013). scRNA-seq enables systematic comparison of cellular composition across species by leveraging gene orthologues to compare gene expression programs rather than individual markers (Marioni and Arendt, 2017). To exploit this opportunity, we next compared our macaque atlases to those previously collected from mice (Macosko et al., 2015; Shekhar et al., 2016), the currently most-studied model. To this end, we adapted the multi-class classification framework used for fovea-periphery comparisons to relate mouse and macaque retinal cell types.

As expected from the conserved retinal architecture, there was a 1:1 match between cell classes in the two species (**Figure S5A**). In addition, there was also a tight correspondence between mouse and macaque PR, HC, BC and AC types. Mice are dichromats, with only M and S cone opsins, and many cones express both opsins (Euler et al., 2014). Mouse M and S cones most closely resembled macaque M/L and S cones, respectively (**Figures 5A, S5B**). Mouse HCs were more transcriptionally similar to macaque H1, which they resemble morphologically, than to H2 (Wassle et al., 2000) (**Figures 5B, S5C,D, Table S2**). For ACs, there was a striking correspondence between macaque and mouse, based on comparison to our previously reported (but underpowered) estimate of 21 molecularly distinct mouse AC clusters (Macosko et al., 2015) (**Figure 5C**). For example, GABAergic and glycinergic types corresponded, and there were clear macaque equivalents of well-studied mouse types such as starburst, AII, glutamatergic (VGluT3+), catecholaminergic, VIP-positive and SEG ACs (**Figures 5D, S5E** and **Table S2**). A more comprehensive ongoing study has revealed >60 mouse AC types, and increased the 1:1 matches with macaque (W.Y., K.S., A.R., and J.R.S. in preparation).

**Figure 5:**
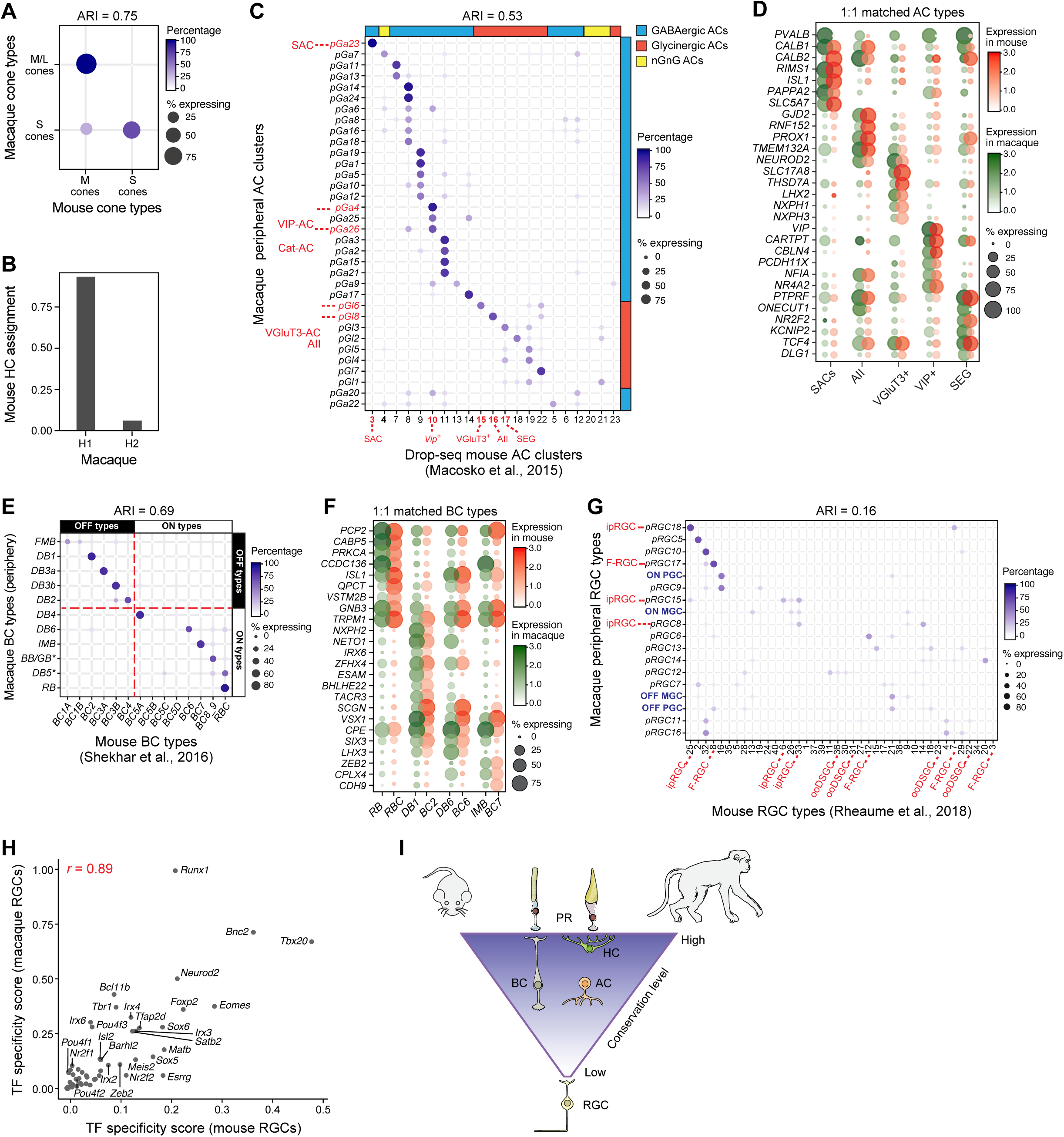
Conservation of retinal cell types between mouse and macaque. A. Transcriptional correspondence between macaque (rows) and mouse cone types identified in (Macosko et al., 2015) (columns). Only mouse cones expressing M- or S-opsin but not both were used for this comparison. B. Supervised classification shows that HCs are more closely related to macaque H1 than to H2 cells. C. Transcriptional correspondence between macaque peripheral AC clusters and mouse AC clusters from (Macosko et al., 2015). The 1:1 mapping of multiple macaque AC clusters reflects the incomplete resolution of AC types in published mouse data. Known types (indicated in red) that map 1:1 are indicated. D. Example of orthologous gene expression patterns in matched macaque-mouse AC types E. Transcriptional correspondence between macaque peripheral BC types and mouse BC types identified in (Shekhar et al., 2016). 9 out of 11 macaque BC types map preferentially to a single mouse type. Crossed red lines highlight correspondence of OFF and ON groups in both species. F. Example of orthologous gene expression patterns in matched macaque-mouse BC types. G. Transcriptional correspondence between macaque peripheral RGC clusters and mouse RGC clusters from (Rheaume et al., 2018). Select mouse and macaque types are indicated. H. Despite poor 1:1 correspondence of RGC types across the two species (**G**), transcription factors that exhibit restricted expression among subgroups are similar, based on high Pearson correlation coefficient (0.89) of their specificity scores (see **Methods**) between macaque and mouse. I. Schematic showing the decreasing molecular conservation of types within cell classes from the outer to the inner retina. For panels A, C, E, G null ARI values (mean ± SD) for a random model are PRs: 1.7 × 10^−4^ ± 9 × 10^−3^, ACs: 2 × 10^−5^ ± 10^−3^, BCs: 3 × 10^−4^ ± 9 × 10^−4^, and RGCs: 2 × 10^−6^ ± 7 × 10^−5^

For BCs, 9 peripheral macaque types mapped preferentially to mouse types with a similar light response (ON vs. OFF, **Figure 5E**; 8 were 1:1), and expression patterns of type-enriched orthologues were conserved between the two species (**Figure 5F**). As noted below, we were unable to identify murine equivalents of MGCs or PGCs, the most abundant primate RGCs. However, the BCs that selectively innervate MGCs (FMB and IMB) correspond to mouse BC1 and 7; and the BCs that provide strongest innervation to PGCs (DB2, 3a and 4) (Tsukamoto and Omi, 2015, 2016) correspond 1:1 to mouse BC4, 3a and 5a. It will be interesting to ask whether postsynaptic targets of these BC types share features with MGCs and PGCs.

### Dramatic divergence in expression programs between macaque and mouse RGC types

In contrast to the correspondence between mouse and macaque BCs and ACs, RGC types differ greatly in both number (>40 in mice (Baden et al., 2016; Bae et al., 2018) vs. ~ 20 in macaque) and distribution (no single mouse type accounts for >10% of all RGCs whereas MGCs account for >50% of macaque RGCs). Moreover, using a recently published scRNA-seq atlas of early post-natal mouse RGCs (Rheaume et al., 2018), we found only a few clear matches between the species (ARI = 0.16), one being the evolutionarily ancient melanopsin-positive intrinsically photosensitive RGC (Do and Yau, 2010) (Figures 5G,I); macaque RGC types exhibit a similar lack of correspondence with adult mouse RGC types (K.S., W.Y., A.R. and J.R.S, *in preparation*). Notably, we found no clear mouse equivalents of macaque ON MGCs, OFF MGCs or OFF PGCs; there may be a mouse equivalent of ON PGCs (**Figure 5G**), but it has not been characterized.

Nonetheless, the RGC comparison did reveal similar patterns of type-enriched transcription factor expression across macaque and mouse RGC clusters. For example, *Tbr1, Eomes, Satb2, Tbx20 and Foxp2*, several of which regulate type-specific features of mouse RGCs (Liu et al., 2018; Mao et al., 2014; Peng et al., 2017), are also expressed by restricted macaque types (**Figure 5H**). Among them, *Satb2* is expressed by bistratified RGCs in both species (**Figure 2I**) (Peng et al., 2017). Moreover, patterns of transcription factor co-expression were similar in both species (**Figure S5F**). For example, *Eomes* and *Meis2* mark largely non-overlapping sets of clusters whereas *Neurod2* and *Satb2* are coexpressed. This suggests a model wherein subfamilies of RGC types in both species are specified by TF codes, but with substantial divergence of the regulatory targets of those codes, accompanied by an abundance of MGCs and lesser diversity in macaques.

### Conservation of cell types and markers across primates

A main value of studies in non-human primates is that their properties are likely to be shared with humans. We asked whether this is true of retina. To this end, we analyzed retinas of humans and another non-human primate, the common marmoset (*Callithrix jacchus*), which is increasingly used because of its small size and genetic accessibility (Mitchell et al., 2014).

We found strong conservation of key molecular and cellular features across macaque, human and marmoset retinas. For example, markers of MGCs, PGCs, S cones and HC types were conserved in marmoset (**Figures 6A, S6A,B**) and the laminar segregation of foveal OFF and ON RGCs was apparent in both marmoset and human (**Figures 6B, S6D**). Distinctions between corresponding peripheral and foveal cell types were also conserved, such as the enrichment of *GNGT1* in marmoset foveal cones (**Figure S6C**) and enrichment of *CYP26A1* and *SPP1* in foveal and peripheral MGs respectively in both humans and marmosets (**Figures 6C,D and S6E, 7E**). Moreover, profiling 2,383 BCs from human peripheral retina, we found close correspondence (ARI = 0.74) between human and macaque types (**Figures 6E,F, S6F-H**), extending the conservation of macaque BC types from mouse to humans. Together with previous morphological studies (Haverkamp et al., 2003; Liao et al., 2016), these results reveal strong conservation of cell types among primates.

**Figure 6:**
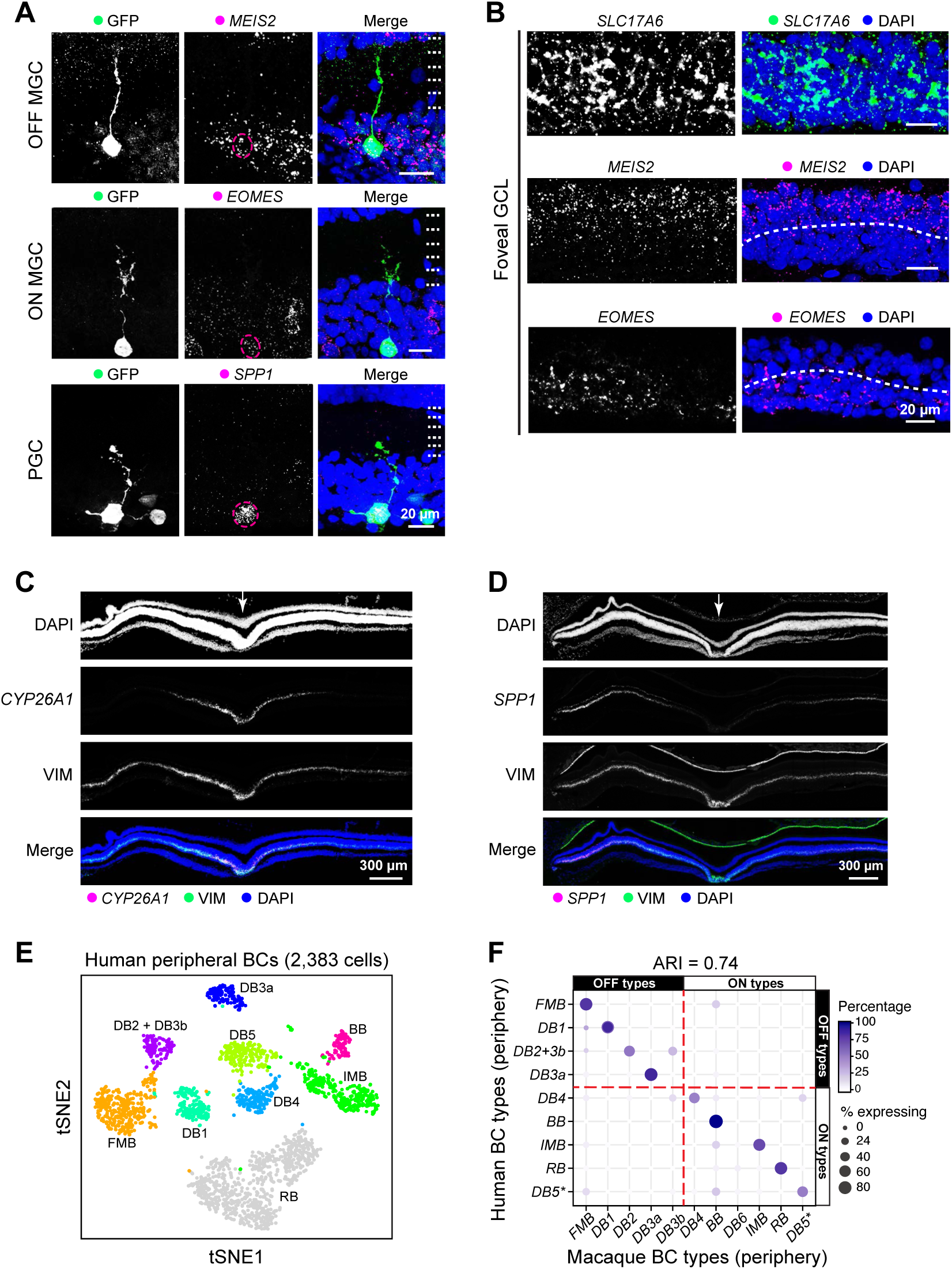
Conservation in marmosets and humans. A. Biolistic labeling combined with FISH shows *MEIS2+* OFF-MGs (top), *EOMES+* ON-MGs (middle), and *SPP1+* PGCs (bottom) in the marmoset fovea. B. The exclusive expression of *MEIS2* and *EOMES* in the marmoset foveal GCL layer. C, D. *CYP26A1* (C) and *SPP1* (D) are selectively expressed by foveal and peripheral Müller glia respectively in marmoset retinas. Arrows indicate the center of the fovea. E. t-SNE visualization of human peripheral BCs. Representation as in **Figure 1D-I**. Clusters are labeled based on their transcriptional similarity to macaque BC types (panel F). F. Transcriptional correspondence between human peripheral BC clusters (rows) and macaque peripheral BC types (columns). Representation as in **Figure 3A**. Each human BC cluster is labeled here and in panel E retrospectively based on the macaque BC type that it maximally corresponds to. Arrows indicate the center of fovea. Scale bars, 20μm and 300 μm as indicated. DAPI staining is blue in A-D.

### Cell type specific expression of human retinal disease genes

Finally, encouraged by the correspondence between human and macaque cell types, we assessed the cellular distribution of genes that have been implicated in 6 groups of human ocular diseases associated with disabling vision loss: retinitis pigmentosa, cone-rod dystrophy, diabetic macular edema and retinopathy, congenital stationary night blindness, glaucoma (including primary open angle and primary angle closure types), and both wet and dry forms of age-related macular degeneration. We focused on ~200 genes for which association has been demonstrated by rare highly penetrant mutations or by GWAS studies (Farrar et al., 2017; Graham et al., 2018; MacGregor et al., 2018; Wiggs and Pasquale, 2017; Zeitz et al., 2015).

For each gene we first calculated an enrichment score for each macaque foveal and peripheral cell class. Aggregating these scores by human retinal disease groups (**Figure 7A**) shows that, in general, expression was highest in cell classes primarily affected by the diseases: PRs for retinitis pigmentosa and rod-cone dystrophy; PRs and BCs for congenital stationary night blindness; RGCs for glaucoma; and non-neuronal cells for diabetic macular edema and retinopathy. As expected, many genes implicated in macular degeneration were expressed in PRs and non-neuronal cells, but several were also highly expressed in other cell classes, especially ACs, a class whose role in disease etiology and progression remains to be explored.

**Figure 7:**
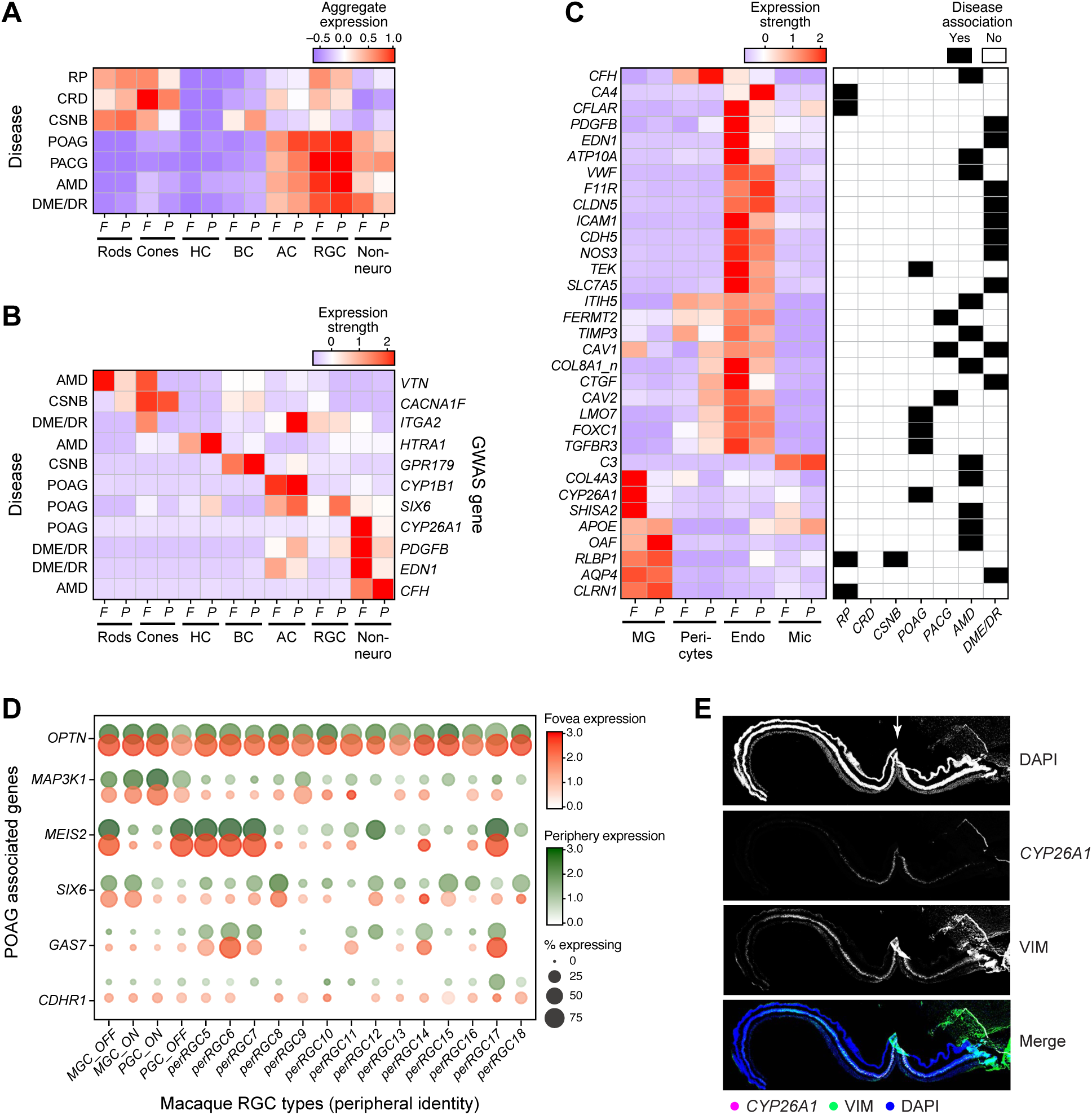
Cell-type and region-specific expression patterns of human retinal disease associated genes. A. Aggregated expression of disease-associated genes in foveal and peripheral cell classes. Disease groups are RP – retinitis pigmentosa; CRD – cone-rod dystrophy; CSNB – congenital stationary night blindness; POAG – primary open angle glaucoma; PACG – primary angle closure glaucoma; AMD – age-related macular degeneration, and DME/DR – diabetic macular edema and retinopathy. B. Expression patterns of a subset of retinal-disease associated genes by cell class (columns), as in panel H. The primary disease associated with each gene is indicated on the right. See **Figure S7** for full list. C. Expression patterns of specific retinal-disease associated genes (rows) by non-neuronal types in the fovea and the periphery (columns). D. Expression patterns of glaucoma associated genes in RGC types in the fovea and the periphery. Foveal RGCs are grouped with their peripheral counterparts. E. The POAG susceptible gene, *CYP26A1*, shows its specific expression in human foveal müller glia. Arrows indicate the center of fovea. Scale bars, 300 μm.

Patterns of expression for several genes are noteworthy. At least three genes associated with diabetic macular edema which, as the name implies, selectively affects the macula, were expressed at higher levels in certain foveal cell types than their peripheral counterparts, such as *PDGFB* and *EDN1* in endothelial cells (**Figure 7C**) (Graham et al., 2018). Likewise, the macular degeneration susceptibility gene *VTN* (Fritsche et al., 2016) was enriched in foveal compared to peripheral rods and cones (**Figure S7**). *HTRA1*, a major susceptibility gene for macular degeneration (Fritsche et al., 2016), which has been implicated in dysfunction of retinal pigment epithelium (anatomically distinct from neural retina and not represented in our dataset) is also expressed at high levels in macaque HCs, but not in mouse HCs. Glaucoma associated genes *CYP1B1* and *CYP26A1* were enriched in a GABAergic AC type and foveal MGs, respectively (**Figure 7B,E and S7**). Several other glaucoma-associated genes were enriched in specific RGC subsets: for example, *MAP3K1* by MGCs and PGCs; *SIX6* by MGCs; and *GAS7* in multiple non-MGC and PGC types (**Figure 7D**). These region- and cell type-selective expression patterns can inform future analyses of genetic variants associated with disease risk. Interestingly, expression patterns of majority of human retinal-disease genes were not conserved in mice, highlighting the limitations of using a mouse atlas as a reference (**Figure S7**).

## DISCUSSION

By profiling >165,000 single-cells, we generated an atlas of molecularly defined cell types in the primate retina. The number of cells needed to generate a comprehensive atlas depends on multiple factors, including the number of types represented in the tissue and the extent to which types differ from each other (Shekhar et al., 2016). In our case, we were able to detect very rare cell types – for example ipRGCs, comprising <0.002% of all retinal cells (Liao et al., 2016). Reasonably complete compendia of cell types from comparably complex tissues will likely require at least similar cell numbers. Using this atlas as a foundation, we then explored regional differences within a tissue and examined evolutionary specializations across species. We undertook two approaches to achieve this: comparing the cellular compositions of fovea and peripheral retina of macaques; and comparing retinal cell types between macaque (which have a fovea) and mice (which do not).

Comparing the fovea and the peripheral retina revealed an overall similar “parts list” but specializations in cellular proportions and cell-intrinsic expression programs of the fovea. Our results also suggest that the compositional differences between the fovea and the periphery are higher at the sensory and preprocessing layers (PRs and BCs) than for downstream processors and feature detectors (ACs and RGCs). A few cell types are unique to or highly enriched in each region, and these may underlie certain functional differences between them. The OFFx bipolar is particularly interesting in this regard, and the absence of some foveal GABAergic AC types may contribute to the low level of foveal inhibition (Sinha et al., 2017). Nevertheless, despite these rare unique types, our results suggest that the dramatic specializations of foveal circuitry and function arise from regional differences in proportion and gene expression within shared types rather than the presence of region-specific types. Some of the differentially expressed genes, such as *GNGT1* and subunits of the GABA receptor, may underlie the differences in the visual responses mediated by the foveal and peripheral circuitry. These differences might arise from developmental programs that specify the fovea (Bringmann et al., 2018; da Silva and Cepko, 2017; Hendrickson et al., 2012; Hoshino et al., 2017). Future studies will focus on elucidating the functions of such region-specific genes to understand signaling processes in the fovea.

By comparing the macaque and the mouse retinal cell taxonomies, we provide compelling evidence for orthology across cell classes, with the dramatic exception of RGCs. The known evolutionary conservation of the fundamental retinal plan extends from cell classes to cell types, with strong conservation for PRs and interneurons (HCs, BCs and ACs). However, this conservation is tenuous for RGCs at the level of gene expression patterns, number and frequency distribution of types. Most notably, MGCs account for most retinal output neurons in primate but lack clear counterparts in mouse. These differences suggest that the outer retina may comprise a conserved set of information processors, with adaptation to species-specific visual processing requirements beginning at the level of RGCs, and then continuing centrally.

Interestingly, despite dramatic differences between macaque and mouse RGCs, there may be an underlying shared genetic regulatory program that specifies RGC diversity, as evidenced by the conserved patterns of transcription factor expression and co-expression restricted to small sets of RGC types. These factors may specify groups that diversify differentially in the two species. Together with recent comparative transcriptomic studies of other parts of the nervous system (La Manno et al., 2016), this raises the possibility that cross-species differences in cell type diversity in homologous regions can be explained by a rewiring of the downstream programs specified by a conserved set of transcription factors.

Finally, our study provides both technical and conceptual foundations for investigations aiming to understand cellular mechanisms underlying blinding retinal diseases. By assessing expression of nearly 200 genes implicated in human retinal diseases, we found that many are selectively expressed in particular retinal cell classes, in particular types within classes, and in foveal or peripheral cohorts on particular types. For example, *HTRA1*, one of the two major risk genes for macular degeneration, is expressed predominantly by HCs among retinal neurons; its role, if any, in these cells merits investigation. Likewise, *PDGFB* and *EDN1*, implicated in diabetic macular edema, are expressed at significantly higher levels by foveal than peripheral endothelial cells, suggesting a basis for the increased susceptibility of the macula in this disease. Interestingly, several of the patterns we document in macaque are not conserved in mice -for example, HTRA1 expression in HCs-emphasizing the value of using primate models for investigating human retinal dysfunction.

## ACKNOWLEDGMENTS

We thank R. Born, A. Liu and J. Wiggs for advice; and L. Gaffney, A. Hupalowska and E. Martersteck for assistance. This work was supported by grants from the NIH (MH105960, EY025840, EY028633, EY025555 and EY028625) and the BrightFocus Foundation (M2014055).

## AUTHOR CONTRIBUTIONS

Y.-R.P., K.S., A. R., and J.R.S conceived the study and wrote the manuscript with input from all authors. K.S., A.S. and W.Y. performed bioinformatic analysis, with guidance from A. R.. Y-R.P. and D.H. performed molecular and histological experiments, with guidance from M. T. H. D and J. R. S.. T.v.T. and G.S.B. procured and prepared tissue. M.T.H.D. provided guidance on study design and analysis.

## METHODS

### Tissue Procurement

Non-human primate and human retinas were obtained and used in accordance with the guidelines for the care and use of animals and human subjects at Harvard University and Boston Children’s Hospital, and supplying institutions. All the procedures on non-human primates were approved by the Institutional Animal Care and Use Committees. Acquisition and use of human tissue was approved by the Human Study Subject Committees (DFCI Protocol Number: 13-416 and MEE - NHSR Protocol Number 18-034H).

Eyes from male macaques (*Macaca fascicularis*, 3-9 years of age) were kindly provided by institutions including Biomere and Massachusetts General Hospital. Eyes were collected either pre-mortem under deep anesthesia or ≤ 45 min post-mortem. In some cases, whole globes were immediately placed in ice-cold Ames solution (Sigma-Aldrich; equilibrated with 95% O2/5% CO2 for all use), where they were stored before experimentation. In others, a rapid hemisection was performed to remove the vitreous and the anterior chamber, and the posterior eyecup was immersed in room-temperature Ames. Retinas were then dissected free and stored in ice-cold Ames solution. Experiments commenced within 8 hours. For access to macaque tissue, including the samples used in this study, we thank V. Belov, C. Cetrulo, D. Guberski, A. Hall, P. Kovalenko, A. LaRochelle, J. Madsen, M. Nedelman, M. Papisov, and S. Smith.

Eyes from male and female common marmosets (*Callithrix jacchus*, 2-10 years of age) were generously provided by the McGovern Institute for Brain Research (Massachusetts Institute of Technology). Animals were perfused with 4% paraformaldehyde (PFA) under deep anesthesia and post-fixed for 1 hr in 4% PFA. Eyes were then collected and stored in ice-cold Ames solution. We are gratefully to G. Feng, Q. Zhang and C. Wu for access to this tissue.

Macaque and marmoset eyes were collected from animals that had reached the end of unrelated studies at supplying institutions. No ocular or visual abnormalities were noted. Data presented in this manuscript did not covary with any treatment that had been applied to the animals.

The human eye used for sequencing and *in situ* hybridization was collected ~6 hours postmortem from a 74 year-old male. The whole globe was immediately transported back to the lab in a humid chamber. Hemisection was performed to remove the anterior chamber and the posterior pole was recovered in Ames equilibrated with 95% O_2_/5% CO_2_ before further dissection and dissociation. The donor was confirmed to have no history or clinical evidence of ocular disease or intraocular surgery We are grateful to Dr. Dejan and the Rapid Autopsy Program, Susan Eid Tumor Heterogeneity Initiative, Massachusetts General Hospital for expeditious access to this material, enabling recovery of high quality RNA for sequencing.

Other human eyes used for *in situ* hybridization were provided by the Lions VisionGift (Portland, OR) and the National Disease Research Interchange (NDRI; Philadelphia, PA). They were collected <11 hr postmortem from a 28 year-old male and a 69 year-old female, hemisected, and fixed overnight in ice-cold 4% PFA following removal of the cornea. No ocular disease was reported in these donors.

### RNA-sequencing

#### Single Cell Isolation

0.5-1.5 mm diameter foveal tissues centering on the foveal pit were dissected out from four macaque eyes. Foveal samples were digested with 200 units papain (Worthington, LS003126). Foveae M1 and M2 were digested at 37°C; foveae M3 and M4 were digested at 22°C. Following digestion, retinas were dissociated and triturated into single cell suspensions with 0.04% bovine serum albumin (BSA) in Ames solution. Peripheral retinal pieces were dissected and pooled from all quadrants of the retina. Single cell suspensions were dissociated at 37°C as described for fovea. Dissociated cells were incubated with CD90 microbeads (Miltenyi Biotec, 130-096-253) to enrich RGCs or with anti-CD73 (BD Biosciences, clone AD2) followed by anti-mouse IgG1 microbeads (Miltenyi Biotec, 130-047-102) to deplete rods. CD90 positive cells or CD73 negative cells were selected via large cell columns through a MiniMACS Separator (Miltenyi Biotec). Single cell suspensions were diluted at a concentration of 500-1800 cells/μL in 0.04% BSA/Ames for loading into 10X Chromium Single Cell A Chips. Human cells were dissociated and treated with anti-CD73 as above.

#### Droplet-Based scRNA-seq

Single cell libraries were prepared using the Chromium 3’ v2 platform (10X Genomics, Pleasanton, CA) following the manufacturer’s protocol. Briefly, single cells were partitioned into Gel beads in EMulsion (GEMs) in the GemCode instrument followed by cell lysis and barcoded reverse transcription of RNA, amplification, shearing and 5’ adaptor and sample index attachment. On average, approximately 10,000 single cells were loaded on each channel and approximately 6,000 cells were recovered. Libraries were sequenced on the Illumina HiSeq 2500 (Paired end reads: Read 1, 26bp, Read 2, 98bp).

#### Viral Transduction

Macaque or marmoset retinas were divided into four quadrants according to cardinal axes with the macula in the temporal region and mounted on Millicell culture inserts (Millipore) with the ganglion cell layer facing up. Retinas were infected with 5 μl of AAV retrograde virus encoding GFP directed by the CAG promoter (Addgene) with a titer ~10e12. Methods were adapted from (Johnson and Martin, 2008; Meyer-Franke et al., 1995). Transduced retinas were cultured in Neurobasal A medium (Fisher 10888-022) supplemented with 0.1 mg/ml penicillin/streptomycin (Sigma G4333), 1 mM L-glutamine (Sigma G7513), B27 (Fisher 17504-044), 0.005 mM forskolin (Sigma F6886), 1 mM sodium pyruvate (Fisher 11360-070), 25 ng/ml BDNF (PeproTech 450-02), and 10 ng/ml CNTF (PeproTech 450-13) for four days. Retinas were then fixed in 4% PFA in phosphate-buffered saline (PBS) for 1hr at 4 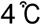 and separated from the culture insert after fixation.

#### Biolistic Transfection

The biolistics procedure was adapted from published protocols (Masri et al., 2016; Morgan and Wong, 2008; Santina and Ou, 2018). Briefly, gold particles (1.6 μm diameter; 12.5 mg; Bio-Rad) were coated with pCMV-GFP DNA plasmid (25 μg). The particles were delivered to retinal cells in whole-mount retinas with GCL facing up using a Helios gene gun (Bio-Rad). The retinal explants were cultured for 3 days followed by fixation as described for virus transduced retinas.

#### Fluorescent In Situ Hybridization

Eyes were fixed in 4% PFA. Marmoset and human eyes were fixed by perfusion and immersion respectively as described under “Tissue Procurement.” For macaque eyes, slits or windows were made in the cornea of eyes collected post-mortem, and the globe was immersion-fixed overnight in ice-cold 4% PFA. Probe templates were generated using cDNA derived from adult macaque, marmoset, or human retinas following RNA extraction and reverse transcription with AzuraQuant™ cDNA Synthesis Kit (Azura, AZ-1995). Antisense probes were generated by PCR using a reverse primer with a T7 sequence adaptor to permit *in vitro* transcription (see Table S4 for primer sequences). DIG rUTP (Roche, 11277073910), DNP rUTP (Perkin Elmer, NEL555001EA), and Fluorescein rUTP (Roche, 11685619910) were used for synthesizing probes for single, double or triple fluorescent *in situ* hybridization (Kolb et al.) experiments. Retinas fixed as above were rinsed with PBS, immersed with 30% sucrose overnight at 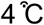, embedded in Tissue Freezing Medium (EMS) and cryosectioned at 20 μm. Details of FISH were described in ref. 12. Briefly, sections were mounted on Superfrost slides (Thermo Scientific), treated with 1.5 μg/ml of proteinase K (NEB, P8107S), and then post-fixed and treated with acetic anhydride for deacetylation. Probe detection was performed with anti-DIG HRP (1:1000), anti-DNP HRP (1:500), anti-Fluorescein (1:1000) followed by tyramide amplification. Detection of protein epitopes was performed following probe detection. Antibodies were diluted in block consisting of 3% donkey serum (Jackson, 017-000-121), and 0.3% Triton-X in PBS at concentrations of 1:500 (anti-GFP), 1:1000 (anti-Calbindin), 1:2000 (anti-PKCα), and 1:100 (anti-CD15) (See below for antibody information).

#### Immunohistochemistry

Tissue was fixed as described above. Immunostaining was performed as described in refs. 12 and 31. Antibodies used were as follows: chick and rabbit anti-GFP (1:500, Abcam; 1:2000, Millipore); mouse anti-Satb2(1:1000, Abcam); goat anti anti-CHX10 (1:300, Santa Cruz); rabbit anti-TFAP2B (1:1000, cell signaling); goat anti-Choline Acetyltransferase (1:500, Millipore); goat anti-VAChT (1:1000, Millipore); rabbit and guinea pig anti-RBPMS (1:1000, Abcam; 1:5000, PhosphoSolutions); rabbit anti-Calbindin (1:2000, Swant); mouse anti-Calretinin (1:5000, Millipore); mouse anti-Gad65/67 (1:1000, Millipore); mouse and rabbit anti-PKCa (1:2000, Abcam; 1:2000, sigma); rabbit anti-Secretagogin (1:10,000; BioVendor); mouse anti-Human CD15 (1:100, BD); rabbit anti-GNGT1 (1:1000, gift from Dr. N. Gautam), mouse anti-7G6 (1:1000, gift from Dr. Peter MacLeish), and chick anti-OPN1M/LW (1:2000, gift from Dr. Jeremy Nathans). Nuclei were labeled with DAPI (1:1000, Invitrogen). Secondary antibodies were conjugated to Alexa Fluor 488, 568, and 647 (Invitrogen) and used at 1:1000. Sections and whole mounts were mounted in ProLong Gold Antifade (Invitrogen).

#### Image Acquisition, Processing and Analysis

Images were acquired on Zeiss LSM 710 confocal microscopes with 405, 488-515, 568, and 647 lasers, processed using Zeiss ZEN software suites, and analyzed using ImageJ (NIH). Images were acquired with 16X, 40X or 63X oil lens at the resolution of 1024×1024 pixels, a step size of 0.5-1.5μm, and 90μm pinhole size. ImageJ (NIH) software was used to generate maximum intensity projections and neuronal dendrites were reconstructed with ImageJ plugin simple neurite tracer. Adobe Photoshop CC was used for adjustments to brightness and contrast.

#### Assembling a Retina-specific Transcriptome for M. fascicularis

We obtained high quality total RNA from macaque retinal tissue (RNA Integrity Number (RIN) score 9.8), and prepared strand-specific libraries using the TruSeq strand-specific Total RNA kit (Illumina Inc.), which was sequenced on the NextSeq 500 system to obtain 45 million 100bp paired end reads. We used StringTie (v1.3.3) (Pertea et al., 2015) to assemble a genome-guided transcriptome, using an available NCBI transcriptome for *M. fascicularis*^1^ (annotation release 101) as an initial guide. Briefly, we mapped RNA-seq reads onto the existing transcriptome in a strand-specific manner using the Hisat2 software (with command line options --dta --rna-strandedness RF), following published guidelines (Pertea et al., 2016). Next, we used StringTie (with command line option --rf) to assemble a new transcriptome based on TruSeq reads. We then reran StringTie (with the command line option merge) to obtain an updated transcriptome annotation, which contained modifications of transcript body definitions from the existing NCBI reference as well as novel transcripts supported by the TruSeq reads. While the modified transcripts retained gene names from the original annotation, the novel transcripts were initially named according to Stringtie’s naming convention (e.g. MSTRG.5141). We call this the NCBI + TruSeq reference in **Figure S1**. There were a few instances where sense and antisense transcripts were predicted for some genes; in these cases we added suffixes “_p” and “_n” against the corresponding gene name to indicate this fact (e.g. *FEZF1_p* and *FEZF1_n*).

To facilitate transcriptional mapping of macaque types to mouse and human types, we used BLAT (Kent, 2002) to associate each macaque transcript with its closest ortholog (reciprocal best matches) in the NCBI mouse (version GRCm38) and human (version GRCh37) transcriptomes, respectively. Encouragingly, this automated procedure matched transcript gene names with similar letter codes (e.g. macaque *TRPM1* with mouse *Trpm1* and human *TRPM1*). We use this mapping to associate each reconstructed transcript with the closest human gene by sequence, in cases where such homology was strong. In some cases, we also found that certain tenuously named loci in the NCBI reference could be associated with homologous mouse and human gene names (e.g. macaque *LOC101864869* could be associated with mouse *Rps27a, LOC102132859* could be associated with *mouse Cyp26a1* etc). For many loci, however, we were unable to associate a human gene and for these cases, we retained the naming convention assigned by StringTie (e.g. MSTRG.5141).

#### Pre-Processing Of 3’ Droplet-based scRNA-Seq Data

Sample demultiplexing, alignment to the NCBI+TruSeq reference, quantification and initial quality control (QC) was performed using the Cell Ranger software (version 2.1.0, 10X Genomics) for each sample (*i.e*., 10X channel) separately. We used the option “--force-cells 6000” in Cell Ranger count to obtain a 36,162 genes x 6,000 cells count matrix each sample to deliberately extract a larger number of cell barcodes in the data, as we found that the automatic estimate of Cell Ranger was too conservative, and was unfavorable for small cell-classes like BCs. Here, 6,000 represented a “loose” upper bound on the number of cells that could be recovered given the density of each the cell suspension loaded onto every channel per the manufacturer’s estimates. We grouped the count matrices separately for the foveal and the peripheral samples to generate a consolidated matrix for each region. Only cells which expressed >500 genes were retained for further analysis. Further pruning of cells (low quality cells, doublets) was done for each class separately (see below).

##### Separation of Major Cell Classes

Both the foveal and the peripheral datasets consisted of multiple biological replicates (**Figs. S1G-I**). Moreover, depending on the enrichment method, each sample contained widely different distributions of the main cell classes. For example, the foveal samples typically comprised of ~30% RGCs, ~28% BCs, ~8% ACs (**Figure S1H**), the peripheral CD90+ samples contained ~28% RGCs, 0% BCs, ~50% ACs, and the peripheral CD73-samples contained <1% RGCs, ~30% BCs and ~10% ACs (**Figure S1I**). Moreover, we anticipated that the differences in proportions also extends to types within a class (**Figure 3**), and that foveal and peripheral types likely had molecular differences (**Figure 4**). This made conventional batch correction using methods such as SVA (Leek et al., 2012) and ComBat (Johnson et al., 2007) difficult, because they make strict distributional assumptions that are violated here. Furthermore, covariates that can be used to counteract these assumptions are not known a priori. We also attempted to perform global batch correction using recently published methods that use Canonical Correlation Analysis (CCA; (Butler et al., 2018)) or Mutual Nearest Neighbour (MNN;(Haghverdi et al., 2018)) matching, but in many cases these did not completely remove batch effects, collapsed distinct types with known molecular markers, or appeared to have varying impact on different cell classes.

To make batch correction more tractable and to avoid extensive biases in the initial stages for the reasons stated above, we first analyzed the foveal and the peripheral datasets separately to stratify the cells by their class. We followed our earlier PCA + Louvain-Jaccard graph-clustering pipeline (Shekhar et al., 2016) without batch correction to obtain a set of transcriptionally distinct clusters. At this stage, we deliberately set parameters to “overcluster” the data to avoid combining distinct cell classes. Next, we scored each cell for gene signatures of well-known retinal cell classes classes - Rods, Cones, Bipolar Cells (BC), Horizontal Cells (HC), Amacrine Cells (AC), Retinal Ganglion Cells (RGC), and non-neuronal cells such as Müller Glia (MG) and others (**Table S1**). Classes with shared markers often scored similarly, but with one higher than the other (e.g. *TFAP2A/2B* and *ONECUT1/2* are expressed in ACs and HCs, with *TFAP2A/2B* being higher in the former and *ONECUT1/2* being higher in the latter). In such cases, we combined these classes and analyzed them as a group in downstream pipelines. Thus, we separated the foveal and peripheral datasets into 5 groups comprising - (i) Rods and Cones, (ii) Amacrine Cells (Fritsche et al.) and Horizontal Cells (HCs), (iii) Bipolar Cells (BCs), (iv) Retinal Ganglion Cells (RGCs), and (v) Others consisting predominantly of Müller Glia, but also pericytes, endothelial cells and microglia (**Figure. 1C**). Astrocytes are absent from the fovea and are largely confined to the optic fiber layer in the periphery (23). None were recovered from either region in our samples. Encouragingly, cells of the same class were more highly correlated with each other than cells of other classes, supporting this separation (**Figure S1F**).

Although clusters at this stage exhibited batch effects, ~90% of clusters could be unequivocally assigned to a class. The other clusters comprised low quality cells and “doublets.” These were often characterized by overall low gene count that was tightly distributed around the minimum cutoff value (~500-700 genes/cell), the mixed expression of more than one class-specific scores albeit at lower levels than “pure” clusters, and the absence of expression of non-class markers that were specific. Moreover, these cells typically clustered proximally on the dendrogram or tSNE visualization to another large, bonafide cluster, as observed previously (Shekhar et al., 2016). To avoid discarding cells erroneously, we analyzed these clusters separately (not shown), and confirmed that they were doublets and low-quality cells. Because the excluded cells were heavily biased towards low gene-count cells, we could have excluded them *ab initio* by setting a more stringent threshold, but this would have risked excluding genuinely small, but intact single-cell libraries.

#### Dimensionality Reduction, Clustering and Visualization For Each Foveal and Peripheral Group

Following the initial clustering step, we analyzed the 6 major classes of cells - RGCs, BCs, ACs, HCs, PRs, and non-neuronal cells for fovea and periphery separately (**Figures 1D-I**). HCs and ACs were separated for this analysis. We employed the following computational steps to cluster each of these groups as described below.

1. <u>Normalization</u>: Expression values *E_i,j_* for gene *i* in cell *j* were calculated following (Shekhar et al., 2016). Briefly, the Cell Ranger reported UMI (i.e. transcript) count value for each gene *i* in each cell *j* was divided by the sum of the total UMI counts in cell *j* to normalize for differences in library size, and then multiplied by *M*, the median UMI counts for all cells within the group, resulting in Transcripts-per-median (*TPM_i,j_*) values. *E_i,j_* was then calculated as log(*TPM_i,j_* + 1)
2. <u>Identification of highly variable genes (HVGs)</u>: We first calculated the mean (*μ*) and coefficient of variation (*CV*) of transcript counts for each gene in the data. We then computed for each gene, the “excess CV” (or eCV) by subtracting the observed CV from a null model of CV vs μ. This null model is based on a Poisson-Gamma mixture, which was shown to accurately model the null CV vs μ relationships for data containing UMIs for a wide range of 3’-biased protocols (Pandey et al., 2018). We calculated the mean and standard deviation (sd) of the eCV values, and selected genes (HVGs) that had eCV > mean + 0.7*sd. This typically selected 700-2500 HVGs for each group.
3. <u>Batch correction</u>: We restricted the expression matrix *E_i,j_* to HVGs and used a linear regression model (adapted from the source code of the ‘RegressOut’ function of the R package ‘Seurat’) to remove correlations of expression values associated with three covariates – the animal of origin (**Figure S1G**), the total number of genes observed per cell, and the expression strength of ribosomal genes within each cell. The choice of covariates was guided by running initial tests on PRs, BCs and RGCs where some prior knowledge of the underlying types exists. We tested different combinations of covariates, and settled on a combination that led to the alignment of known types across replicates (e.g., cones or IMBs), and used this scheme consistently for all cell group analyses. Here, we also tested the CCA-based (Butler et al., 2018) and MNN-based (Haghverdi et al., 2018) batch correction strategies, but found that the performance of these methods depended on the choice of certain tunable parameters, which were different for different groups. The linear regression model, on the other hand, did not contain additional parameters outside of the choice of covariates, and was computationally much faster than the other two methods. In a few cases, some residual batch effects still remained, which we corrected for in a supervised manner (see below). For BCs and RGCs, two groups extensively validated in this study, we additionally confirmed the absence of any artifacts by re-clustering single batches separately and verifying that all the abundant clusters were preserved. We refer to the corrected expression matrix as 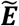. For each of the HVGs, we compute the Pearson correlation coefficient between the values in ***E*** and those in 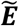. Encouragingly, most genes had a similar correlation coefficient (r > 0.97), but ~30 genes had a significant reduction in correlation (r < 0.6). We flagged these “batch specific genes” in our differentialexpression analysis of clusters in 5 (see below).
4. Principal component analysis and 2D visualization: We restricted the corrected expression matrix 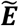 to HVGs, and values were centered and scaled before input to Principal Component Analysis (PCA), which was implemented using the R package ‘irlba’. Statistically significant PCs were estimated using the Tracy-Widom distribution (Kiselev et al., 2017; Shekhar et al., 2016), which we found to be an excellent approximation to exact values computed using the more exact, but computationally intensive, bootstrap permutation test (Shekhar et al., 2016). For visualization purposes only, dimensionality was further reduced to 2D using t-distributed stochastic neighbor embedding (t-SNE). We used a recently developed rapidly accelerated implementation (Linderman, 2017) of the original algorithm (Hinton, 2008) setting the “perplexity” parameter to 30. The resulting t-SNE cell coordinates were used to generate the visualizations in **Figure 1D**-I.
5. <u>Graph-based clustering</u>: To partition the data into clusters of transcriptionally related cells, we used unsupervised clustering based on the Louvain algorithm (Vincent D. Blondel, 2008) with the Jaccard correction (Levine et al., 2015; Shekhar et al., 2016). We first constructed a *k*-nearest neighbor graph (kNN) using the Euclidean distance metric between pairs of cells in the reduced dimension space of the significant PCs, as implemented in the R package ‘RANN’. k=30 was used for all groups, except peripheral HCs where k=15 was used because he low cell count.
6. <u>Cluster refinement</u>: As the Louvain method suffers from a well-known “resolution” limit (Fortunato and Barthelemy, 2007), we refined the set of initial clusters as well as corrected for potential overclustering due to residual batch effects in three steps as follows. *Step 1* and *Step 2* were performed for each of the initial clusters in (5) separately. In *Step 1*, each cluster was sub-clustered using the more sensitive Infomap community detection algorithm (Rosvall and Bergstrom, 2008) on a new *k*-nearest neighbor graph (k = 5*log(cluster size)). However, this often ended up overclustering the data, as noted before (Shekhar et al., 2016). In *Step 2*, the subclusters within each cluster were assessed for differential expression, and iteratively merged as follows. We built a dendrogram on the subclusters (hierarchical agglomerative clustering with Euclidean distance, complete linkage), and tested the closest three pairs of leaves of the dendrogram for differential gene expression using the MAST test (Finak et al., 2015), corresponding to the three pairs of the most transcriptionally similar subclusters. If any pair did not show sufficient differential expression (minimum 5 DE genes enriched both ways, fold-change > 1.4, p-value < 0.001, excluding mitochondrial transcripts and ribosomal protein genes, or “batch genes” defined above), the corresponding subclusters were merged. If any of the three pairs of subclusters were merged, we rebuilt the dendrogram using the new subclusters, and repeated this procedure until convergence. *Step 1* and *Step 2* resulted in the splitting of some of the original Louvain clusters. In *Step 3* we built a dendrogram on the new set of clusters and iteratively merged (as in *Step 2*, except with more stringest parameters) until convergence. During this final dendrogram-based merging round, we elected to merge transcriptionally proximal clusters using a more stringent rule. In addition to the criteria described above (≥ 5 DE genes enriched both ways, fold-change > 1.4, p-value < 0.001, excluding mitochondrial and ribosomal, or batch-specific genes), we required that the two clusters being tested be distinguished by ON/OFF expression (i.e. not just differences in expression levels) of at least one gene in each of them. More specifically, we required that each of the two clusters show at least one gene that was expressed in at least 4-fold higher proportion of cells the other cluster (e.g. ≥80% of cells in cluster 1 and ≤20% of cells in cluster 2, and ≥60% of cells in cluster 2 and ≤15% of cells in cluster 2).
7. <u>Removing doublets and contaminants</u>: Although dimensionality reduction and clustering were performed separately for each cell class after an initial segregation, we noted that the data was not entirely free of contaminants. For example, we found small PR-like or AC-like clusters when analyzing BCs in the fovea or periphery. These were likely doublets as these clusters also expressed BC-markers, and comprised <1-2% of the cells in the dataset. While such clusters appeared more BC-like in the initial stage, when all cell classes were analyzed together, their contaminant-like features became more apparent when analyzed only with pure BCs. These contaminant-like clusters were removed manually, and we verified that the final clusters for each cell-class expressed at most negligible levels of markers of other cell-classes.

#### Distinguishing M vs L Cones

The transcript sequences of *OPN1MW* and *OPN1LW* are extremely similar across Old World Monkeys and humans (Nathans et al., 1986; Onishi et al., 2002). In the crab-eating macaque, there are only 26 single nucleotide polymorphism (SNPs) over 1,095 nucleotides of the full-length cDNA sequences of *OPN1MW* and *OPN1LW*, located between exon 2 and exon 5 (Onishi et al., 2002). This made it difficult to unambiguously assign a short read (98bp) aligning to this locus to either *OPN1MW* and *OPN1LW*, as SNPs could not be distinguished from sequencing errors. Thus, cells in the foveal cone cluster that expressed transcripts aligning to this locus, but not the S-opsin *OPN1SW*, were initially tagged as “M/L cones.” However, these SNPs could be robustly distinguished from sequencing errors when aggregating across *all* the reads aligning to this locus (**Figure S2A**), and their positions were consistent with previous studies (Onishi et al., 2002). For each M/L cone cell, we then counted the *OPN1MW-* and OPN1LW-specific SNPs (**Figure S2B**). ~70% of M/L cones in the data possessed reads that had at least one transcript covering one of the 26 SNPs. Cells with exclusively *OPN1MW-* and *OPN1LW*-specific SNP counts were labeled M and L-cones, respectively, and the remainder were called “mixed” (**Figures S2D,E**).

#### Differential Expression Analysis

Differential expression (DE) tests for a gene between a pair of clusters or between a cluster and the remaining clusters were performed using the ‘MAST’ package (Finak et al., 2015), which implements a two-part ‘hurdle’ model to control for both technical quality and cross-animal variation. P-values were corrected for multiple hypotheses by controlling the false discovery rate (FDR; (Hochberg, 1995)) using the R function ‘p.adjust’.

We sought markers for a cluster by comparing it to other clusters only within the same class for DE genes, since class specific markers are well-defined (**Table S1, Figs. 1C**). To obtain highly specific markers for histological validation, we ranked significantly enriched (>1.5-fold difference, FDR adjusted p < 10^−6^) markers in increasing order of the proportion of background cells where they were detected (**Table S3**). Differential gene expression patterns were visualized either in dotplots (e.g. **Figure 2B**) or box-and-whisker plots (e.g. **Figure 4B**).

To identify fovea or periphery specific DE genes for cell types that had 1:1 matches across the two regions, we collected the corresponding cells from each region and applied MAST as before, while controlling for library size differences. In choosing candidates to pursue for validation, we also avoided weakly enriched genes that could be explained by large differences in abundance of a different cell class in the same region. For example, *RHO* and *GNAT1* were weakly enriched in peripheral RGCs, BCs and ACs, which likely reflects contamination from the predominant rods in the periphery.

#### Transcriptional Mapping of Foveal Types to Peripheral Types within A Class

We compared the consistency between foveal and peripheral clusters for HCs, BCs, ACs and RGCs within each cell class using a multi-class classification approach, as described before (Shekhar et al., 2016). For PRs and non-neuronal cells, the fovea vs periphery mapping was straightforward as the types could be easily identified based on known markers, and their transcriptional identities could be matched between the two regions.

For BCs, RGCs, ACs and HCs, we trained a multi-class classifier using Random Forests (RF;(Breiman, 2001)) as well as the recently published Xgboost algorithm, which uses gradient boosted trees (Tianqi Chen, 2016) using the R packages ‘randomForest’ and ‘xgboost’, respectively. In each case, we trained the classifier on the peripheral cluster labels, using as features the common HVGs identified in the foveal and peripheral datasets as above (cluster-specific markers were not favored in any way). The training was performed on 50% of the cells in peripheral datasets and its performance was tested on the remaining, “held-out” dataset. (The foveal data was not used for training.) For each of the 4 classes (HCs, BCs, ACs and RGCs), the maximum error rate for types in the held-out data was < 1% (not shown), suggesting that the type-specific transcriptional signatures were robust, learnable, and not prone to overfitting.

The trained classifier was then used to assign each foveal cell a peripheral identity, in a manner that was agnostic to its foveal cluster identity. For each cell, we computed the “margin”, i.e the fraction of decision trees that vote for the “winner” type (majority vote). We excluded cells which fell into the following criterion,

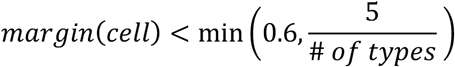

This was done to buffer against potential misclassifications due to a marginal majority.

For all of the cell classes tested, < 4% of cells were excluded due to this criterion. The congruences of the foveal cluster identities and their peripheral matches were visualized as confusion matrices for each class (**Figures 3A-D**). A mapping between a foveal cluster *F* and a peripheral cluster *P* was considered 1:1 if > 85% of cells from *F* mapped to *P*, and if < 5% of cells from every other peripheral cluster mapped to *F*. Results from Xgboost and RF were extremely comparable, although the former was much more computationally efficient. The confusion matrices (**Figures 3A-D**) shown are equally-weighted aggregated results of Xgboost and RF.

To quantify the extent of 1:1 mapping between foveal and peripheral clusters we computed the Adjusted Rand Index (ARI), a measure of the similarity between two data clusterings, between the foveal cluster labels and the Xgboost/RF–assigned cluster labels using the R package “mclust”. The significance of the computed ARI values were evaluated by comparing against “null” ARI values computed by randomly permuting the cluster labels across cells. Mean and standard-deviation of null ARI values were computed by averaging 10,000 trials.

#### Comparison of Cluster Frequencies between the Fovea and the Periphery

We computed frequency differences of 1:1 matched types between the fovea and the periphery, and assessed their significance using a two sample t-test. For foveal types that were a mixture of multiple peripheral types (e.g. fRGC11), we assigned each foveal cell its matched peripheral cluster identity prior to comparison (see previous section). A fovea vs. periphery frequency comparison could not be meaningfully done for ACs, as CD90 non-uniformly labels ACs. We believe that this phenomenon likely explains the poor correlation between foveal and peripheral AC type frequencies (r=0.12); we could therefore not make a fair comparison of AC types between the regions. Interestingly, although the CD73-samples contributed only ~10% of all peripheral ACs, the AC type frequencies in these samples correlated much better with the foveal population (r=0.79; **Figure S4B**), and suggested that a few peripheral types (e.g. VIP+ ACs) were conspicuously underrepresented in the foveal samples (**Figure 3C**).

We quantified the compositional similarity between foveal and peripheral clusterings for each cell class by computing the Jensen-Shannon divergence (JSD), as follows. Let *P =* {*p*_1_*, p*_2_*,…, p_n_*} and *Q =* {*q*_1_*, q,…,q_n_*} be the peripheral and foveal frequencies of *n* clusters of a cell class *C*, each of which sums to unity. Here *n* is the total number of clusters which is a sum of the 1:1 matched clusters, and the clusters unique to either region. Then the JSD between the fovea and the periphery for cell class *C* is computed as,

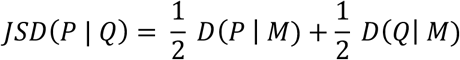

Here, 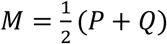, and *D*(*P* | *Q*) is the Kullback-Leibler (KL) divergence between probability distributions *P* and *Q* as follows,

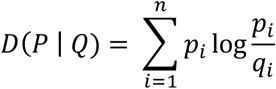

Note that while *D*(*P | Q*) *≠ D*(*Q | P*), *JSD*(*P | Q*) is symmetric in that *JSD*(*P | Q*) = *JSD*(*Q | P*).

In comparing compositional similarity between foveal and peripheral cell classes, we used average frequencies across all replicates for foveal PRs, BCs, HCs, ACs and RGCs, but only BCs, HCs, ACs and RGCs in the periphery. This is because our collection procedure (**Figure S1B**) methods selected against peripheral PRs. We therefore used a published estimate of peripheral PR composition: 95% rods, 4.75% M/L cones and 0.25% S cones (Roorda et al., 2001; Wikler et al., 1990). In addition, we considered only the CD73-samples for ACs because the CD90+ samples differentially excluded some AC types. For peripheral ACs, we only used their composition in the CD73-samples (see above). We expect rods to be underrepresented in our peripheral samples due to our collection procedure.

#### Transcriptional Mapping of Cell Types across Species

*<u>Mouse</u>:* We transcriptionally related macaque cones, HC, BC, AC, and RGC types to their mouse counterparts from earlier publications using the same approach that related foveal to peripheral types, with minor modifications (see below). We conducted the following mappings:

1. 11 macaque peripheral BC clusters (BB/GB* was considered as a single cluster and then subclustered *post hoc*) were mapped to 15 mouse BC types published in our earlier study that identified these types molecularly and matched them to morphology (Shekhar et al., 2016)
2. 34 Macaque peripheral AC clusters were mapped to 21 mouse AC clusters identified in our earlier whole-retina study (Macosko et al., 2015), which represents an incomplete molecular characterization of ACs, estimated to contain >60 types based on morphological (Diamond, 2017) and molecular (JS, WY, KS, AR, and Mallory Laboulaye, *unpublished*) diversity.
3. Macaque peripheral H1 and H2 were transcriptionally related to cells from a single cluster of mouse HCs in our previous whole retina study (Macosko et al., 2015) to determine whether mouse HCs were molecularly more similar to one of the two types.
4. 18 macaque peripheral RGC clusters were mapped to a recently published dataset of postnatal day 5 (P5) mouse RGCs, which identified 40 distinct molecular clusters (Rheaume et al., 2018).
5. As we did not sample peripheral S-cones, we related macaque foveal M/L and S cones to mouse cones from our previous full retina data (Macosko et al., 2015). We computationally separated mouse cones into those that exclusively expressed *Opn1sw* or *Opn1mw* to enrich for pure S-cones and pure M-cones, respectively.

In all cases except HCs, we trained the classifier on the mouse types, and mapped the macaque cells on to these types. In each case 1-5 above, we trained RF/Xgboost classifiers on the training data (mouse for Cones, BCs, ACs, RGCs and macaque for HC) using as features the common set of highly variable genes (HVGs) in both the mouse and the macaque datasets among the 1:1 orthologues. Restricting the features to the common set of HVG orthologues was important for the performance of the classifier; in our initial tests, we found that using HVG orthologues identified only in one of the two species resulted in mappings that were more diffuse, likely due to genes that expressed in one species at high levels but, were completely absent in the other due to transcriptional rewiring. However, we emphasize that no manual curation of the feature set was performed to favor or disfavor certain genes based on their cell-type specific expression. We trained the classifier using 80% of the cells in the relevant mouse dataset, and validated it on the “held-out” cells in that dataset, which showed excellent performance in all cases. Next, we applied the classifier on the test set (macaque Cones, BCs, ACs, RGCs and mouse HCs), and assigned each cell an identity in the other species. As above, cluster identity of cells in the test set was not used to guide the mapping. Mappings of cell types across species were visualized using confusion matrices for Cones, ACs, BCs and RGCs (**Figures 5A,C-E**) and a barplot for HCs (**Figure 5B**). A mapping between a test cluster *C_test_* and a training cluster *C_train_* was considered 1:1 if > 75% of cells from *C_test_* mapped to P, and if < 5% of cells from every other test cluster mapped to *C_train_*.

*<u>Human</u>:* We related human BC clusters to macaque BC clusters using an approach similar to that described above for mouse. We trained a classifier on the macaque peripheral BC data, applied it to map each human BC to a macaque cluster, and visualized the resulting confusion matrix to identify 1:1 matches (**Figure 6F**).

#### Conservation of Transcription Factor (TF) Codes in Mouse and Macaque RGCs

We downloaded a curated list of human and mouse transcription factors from the TFCat Transcription Factor Database Catalog (www.tfcat.ca), CIS-BP database (http://cisbp.ccbr.utoronto.ca/) and (Lambert et al., 2018), and used these to assemble a corresponding list of macaque transcription factors from our orthology list (see above). We filtered this list in two ways. **First**, we restricted the list to those TFs that had 1:1 mouse and macaque orthologs. **Second**, we only considered those TFs that were expressed (> 0 UMIs/cell) in > 25% of the cells in at least one cluster.

For each of the remaining TFs, we computed a TF specificity score (TFSS) among RGC types separately in macaque and mice. Specifically, for a transcription factor *t*, we computed a vector 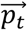 of the fraction of cells expressing *t* in every RGC cluster. For example, 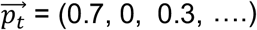 implies that 70% of cells in RGC cluster 1, 0% of cells in cluster 2, 30% in cluster 3, and so on, express transcription factor *t*. We normalized 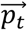 to sum to 1, and computed the Shannon index of the resulting normalized vector,

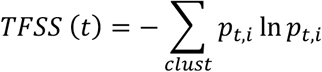

We computed *TFSS_mouse_*(*t*) and *TFSS_macaque_*(*t*) for all filtered transcription factors *t* (n=93), and compared them (**Figure 5G**). Note that constitutively expressed TFs like *Pou4f1/Six6* have low values on the plot, because of their low Shannon indices.

To compute TF co-expression scores and compare across species, we further filtered the 1:1 orthologous TF pairs to only include those TFs that are expressed in at least 15% of clusters in both species. For each species, we then binarized TF expression in RGC clusters as a matrix ***T***, where ***T***(*t, c*) *= 1* if TF *t* is expressed in at least 25% of cells in cluster c. For every TF *t*, we computed its probability of expression in types, *α*(*t*), in any species as,

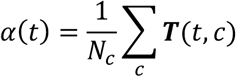

Here, *N_c_* refers to the number of clusters. The observed probability of co-expression of *TFs t_1_* and *t_2_, A*(*t*_1_*,t*_2_), can be computed as,

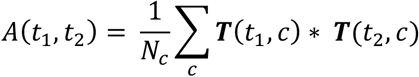

We compute the TF co-expression scores (TFCS) in either species as,

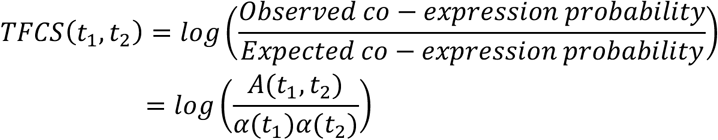

**Figure S5G** compares *TFCS_mouse_*(*t*_1_, *t*_2_) and *TFSS_macaque_*(*t*_1_, *t*_2_).

#### Evaluating Cell-Type Specific Expression of Disease Associated Loci

We assembled lists of genome-wide associated study genes for prominent retinal diseases that cause blindness – primary open angle glaucoma (POAG), primary angle closure glaucoma (PACG), cone-rod dystrophy (CRD), retinitis pigmentosa (Leferink et al.), congenital stationary night blindness (CSNB), age-related macular degeneration (Springelkamp et al.) and diabetic macular edema (DME) from (Farrar et al., 2017; Fritsche et al., 2016; Graham et al., 2018; MacGregor et al., 2018; Wiggs and Pasquale, 2017; Zeitz et al., 2015).

To facilitate 1:1 comparison between the fovea and the periphery, we assigned each foveal cell a peripheral type identity using the RF/Xgboost mapping described above. We only considered those genes that were expressed in at least 20% of cells of at least one cell type, which resulted in 196 genes for further consideration. We note that this does not necessarily rule out the possibility of these genes turning on in specific cell types during disease pathogenesis, just as the *expression* of a gene in a cell type at baseline does not necessarily imply vulnerability in disease.

We visualized expression patterns of the disease genes in our atlas using one of three visualizations:

1. Dotplots, which show cell-type resolution (as in **Figure 7D**),
2. Heatmaps showing relative expression of individual genes in major cell classes (**Figures 7B, S7**), and
3. Heatmap showing aggregated expression of groups of genes corresponding to individual diseases in major cell classes (**Figure 7A**). While 2. and 3. lack cell-type resolution, they do show statistical enrichment of genes and gene groups in specific cell classes, and differences in fovea and periphery, if they exist.

For each of the fovea and periphery, we summarized the expression of all disease genes (see above) across all cell types, by computing two matrices ***E*** and ***F***. For gene *g* and cell-type *c*, ***F***(*g,c*) was the fraction of cells in cell type *c* that had non-zero expression ( > 0 UMIs) of *g*, and ***E***(*g,c*) was the average number of transcripts in cells with non-zero expression. To remove outliers, we capped values in ***E*** to the 99.5^th^ quantile of expression. We then computed a matrix of gene expression scores for all genes across all cell types as the product of these two matrices,

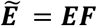

For (2), the we computed the expression strength ***S****(g,C)* of each gene *g* in a cell class *C* (one of Rods, Cones, HCs, BCs, ACs, RGCs, and non-neuronal cells)

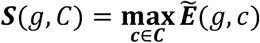

Here, we computed the maximum, rather than the average or aggregate, to highlight specific instances of highly cell-type specific expression of genes in a class. We combined ***S****_fovea_*(*g, C*) and ***S****_periphery_*(*g, C*), and visualized them as a single heatmap, where we z-scored the rows (corresponding to genes) to highlight relative expression strengths 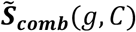.

Specific rows are highlighted in **Figure 6I**. Here, the initial filtering of removing genes that were not expressed in at least 20% of cells in any cell type avoided spurious patterns.

For (3), we subset 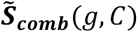 by disease and computed the mean relative expression strengths within each cell type group for each disease. This aggregate expression strength was visualized in **Figure 7A**.

In **Figure S7**, we visualized the relative expression strengths of genes grouped by disease as in **Figure 7B**, but after subdividing non-neuronal cells into MGs, Pericytes, Endothelial cells and Microglia to highlight differential expression among these types.

## SUPPLEMENTAL FIGURE LEGENDS

**Figure S1:**
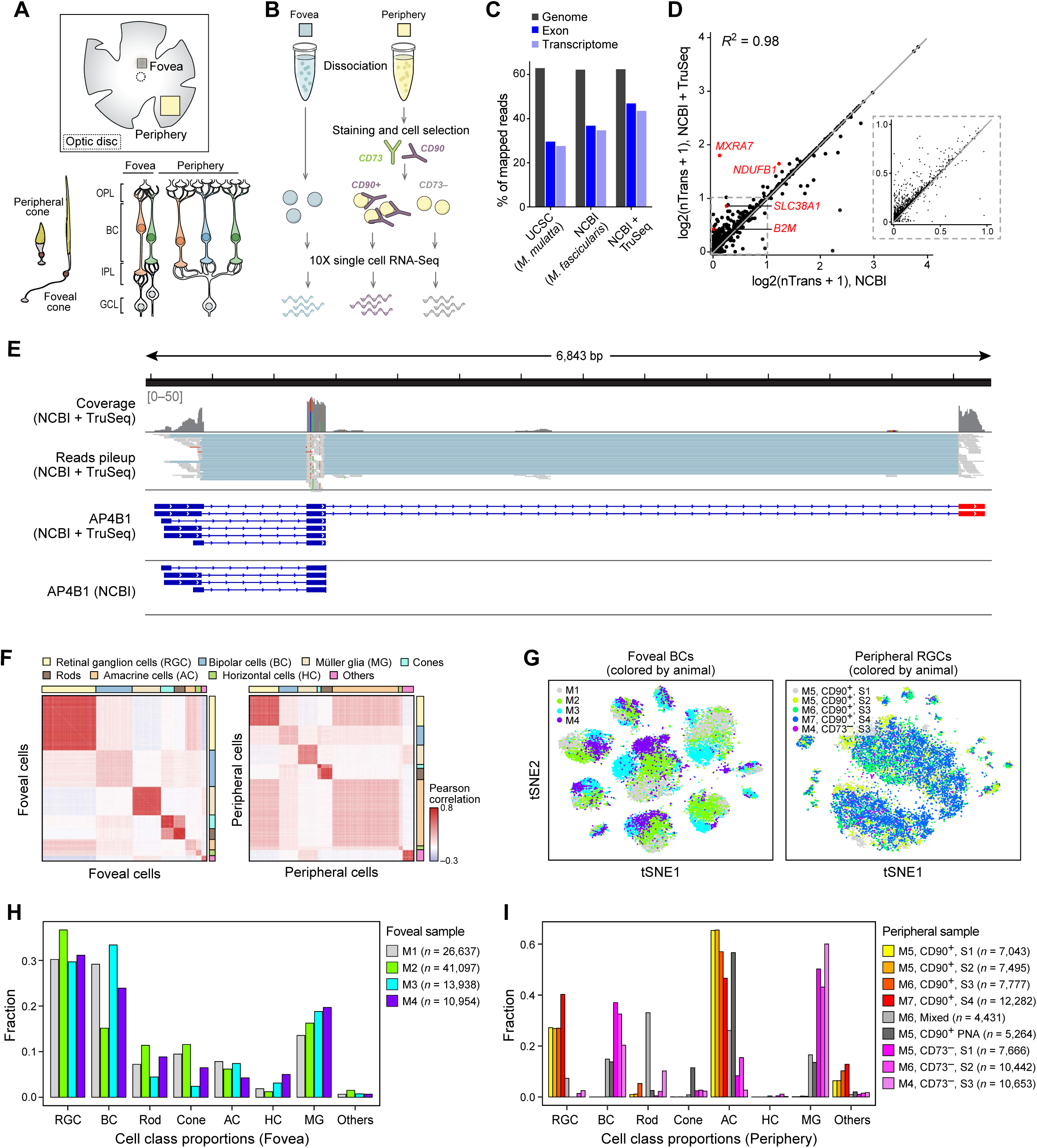
Foveal specializations, experimental design and data quality. A. (Top)The position of fovea and optic disc in flat mount. (Lower right) Sketch of foveal and peripheral cones: foveal cones are longer and more slender than peripheral cones, and bear longer axons. Adapted from (Greeff, 1874). (Lower left) Foveal and peripheral retinal circuits: In the fovea, one cone provides input to two midget BCs, each of which innervates one midget RGC for a cone:midget RGC ratio of 1:2. In the periphery, multiple cones provide input to each midget BC, and multiple midget BCs innervate one midget RGC, for a cone:midget RGC ratio of ≥10:1. Thus the degree of convergence of cones onto interneurons and RGCs differs by an order of magnitude between the fovea and periphery. In addition, peripheral MGCs receive input indirectly (via synapses from rod BCs to AII ACs to cone BCs) from hundreds of rods; whereas foveal MGCs receive little if any rod input. Thus, the total difference in PR:MGC convergence is even greater between regions. Adapted from (Kolb and Marshak, 2003). B. scRNA-seq workflow. Cells were dissociated from <1.5 mm-diameter foveal samples and collected without further processing. Because the peripheral retina is dominated by rod PRs (~80% of cells), we used magnetic columns to deplete rods (CD73+) or enrich RGCs (CD90+). C. Mapping rates of scRNA-seq reads to the genome, exonic and transcriptomic (exonic with splice-junction constraints) regions using three different transcriptomic references – UCSC (University of California Santa Cruz Genome Browser reference for *Macaca mulatta*, a related but different species), NCBI (the publicly available reference for *Macaca fascicularis*) and NCBI + TruSeq (the retina specific transcriptome assembled in this study using the NCBI reference as a starting point). Note that the new transcriptome we used increased the percentage of exonic reads that could be assigned to a gene by ~25% compared to the NCBI reference for *Macaca fascicularis* (from 37% to 47%) and by ~68% compared to the NCBI reference for *Macaca mulatta* (from 28% to 47%). D. Comparison of expression levels (log2(number of transcripts + 1)) of common genes between the NCBI and NCBI + TruSeq references, exhibits a high degree of correlation. A greater number of gene loci exhibit increased expression levels due to higher mapping rate (panel A) as evident by the fact that the majority of points lie above the red line (x=y). This is particularly true for genes expressed at low levels, as shown in the inset. A small number of loci were mapped less well in the improved transcriptome because new transcripts could not be assigned unambiguously to a single gene. E. Example of improved gene-body definition in the assembled transcriptome of *AP4B1* as visualized using the Integrated Genomics Viewer (IGV). The lower panel shows the gene-body definitions for the NCBI and the NCBI +TruSeq references. In this example, the NCBI + TruSeq reference includes a distal 3’ exon that is absent from the NCBI reference. The middle panel show the pile-up of individual reads from a sample 10X run mapped to the expanded locus and the upper panels show the read coverage. Blue shading connects portions of a read that spans a splice junction. The coverage plot shows that a large proportion of reads mapped to 3’ exons present in the NCBI + TruSeq reference (highlighted in red) but absent from the NCBI reference. F. Heatmap of Pearson correlation coefficients between each pair of 92,628 foveal cells (left) or 73,053 peripheral cells (right) (rows and columns) ordered by cell class (annotated as color bars along row and column). “Other” cells are comprised of pericytes, vascular endothelial cells and microglia. G. Examples illustrating the lack of strong batch effects. tSNE visualization of foveal BCs (left, as in **Figure 1F**) and peripheral RGC (right, as in **Figure 1H**), which are now colored by their animal of origin (M1-7), shows the representation of all animals across clusters. H., I. Proportions of major cell classes in foveal (H) and peripheral (I) samples colored by experimental batches. Peripheral samples (I) colored using palettes corresponding to one of four processing methods prior to collection: (1) CD90+: RGCs were enriched on a magnetic column using beads conjugated with antibodies to the RGC class marker, Thy1 (CD90). Cells that did not bind were discarded, and bound cells were eluted and used. (2) CD73-: Rod photoreceptors were depleted by passage over a magnetic column containing beads conjugated with antibodies to the rod-specific marker CD73. Unbound cells were used. (3) Mixed: In this experiment CD90+ cells (magnetic column selection) and non-enriched cells were mixed 1:1 prior to use. (4) CD90+PNA: In this experiment, PNA (Blanks and Johnson, 1984) was added to enrich for cones. Cell numbers in each experimental batch are indicated in parentheses. M4 here corresponds to the same animal. Each experimental batch corresponds to an independent animal, denoted M1-M7. The entire fovea was dissociated, and cells were collected for unbiased sampling of all types. Sequencing batches, typically containing 3000-5000 cells each, are aggregated within experiments as they did not exhibit any appreciable batch effects. Cell numbers in each experimental batch are indicated in parentheses.

**Figure S2:**
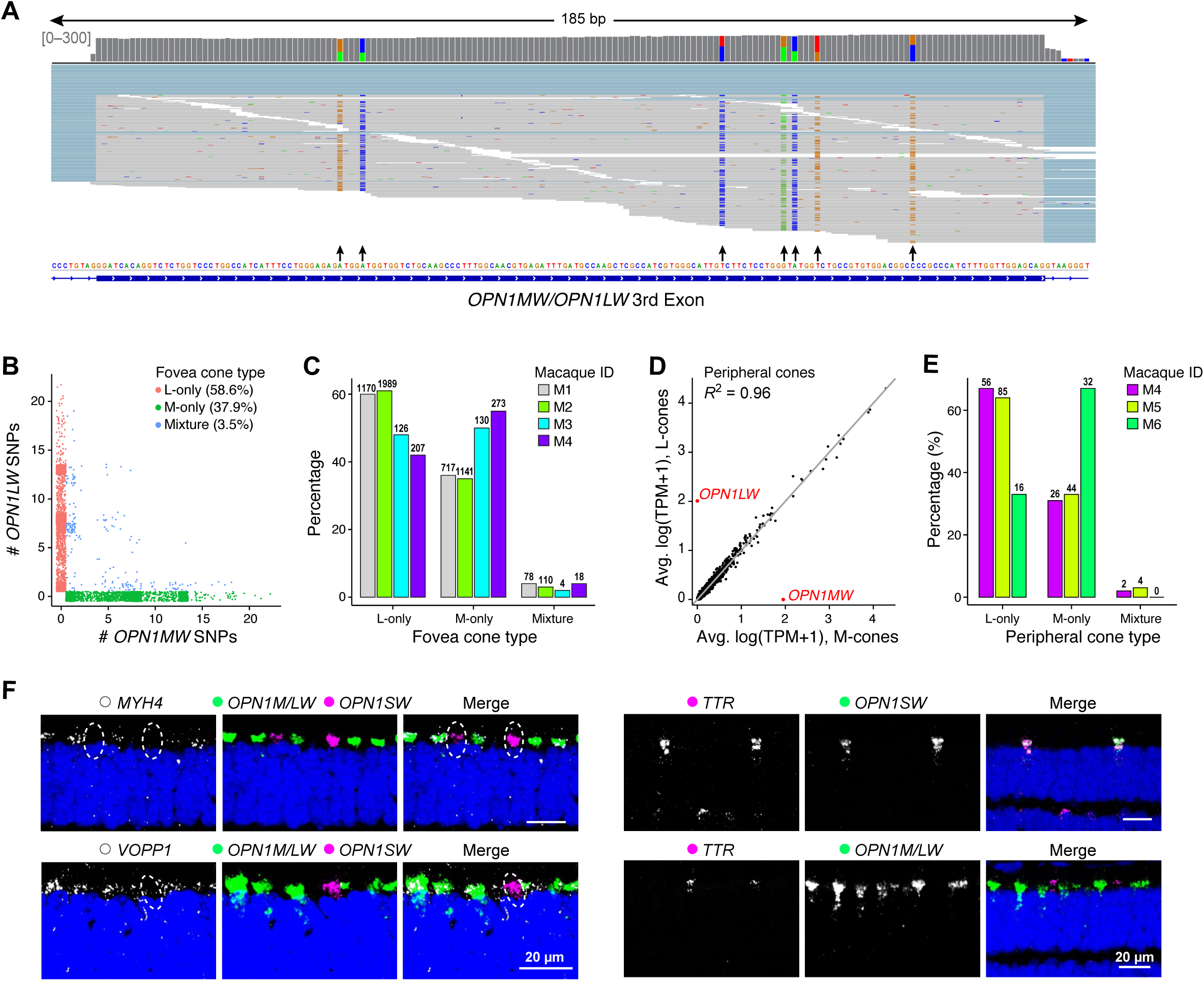
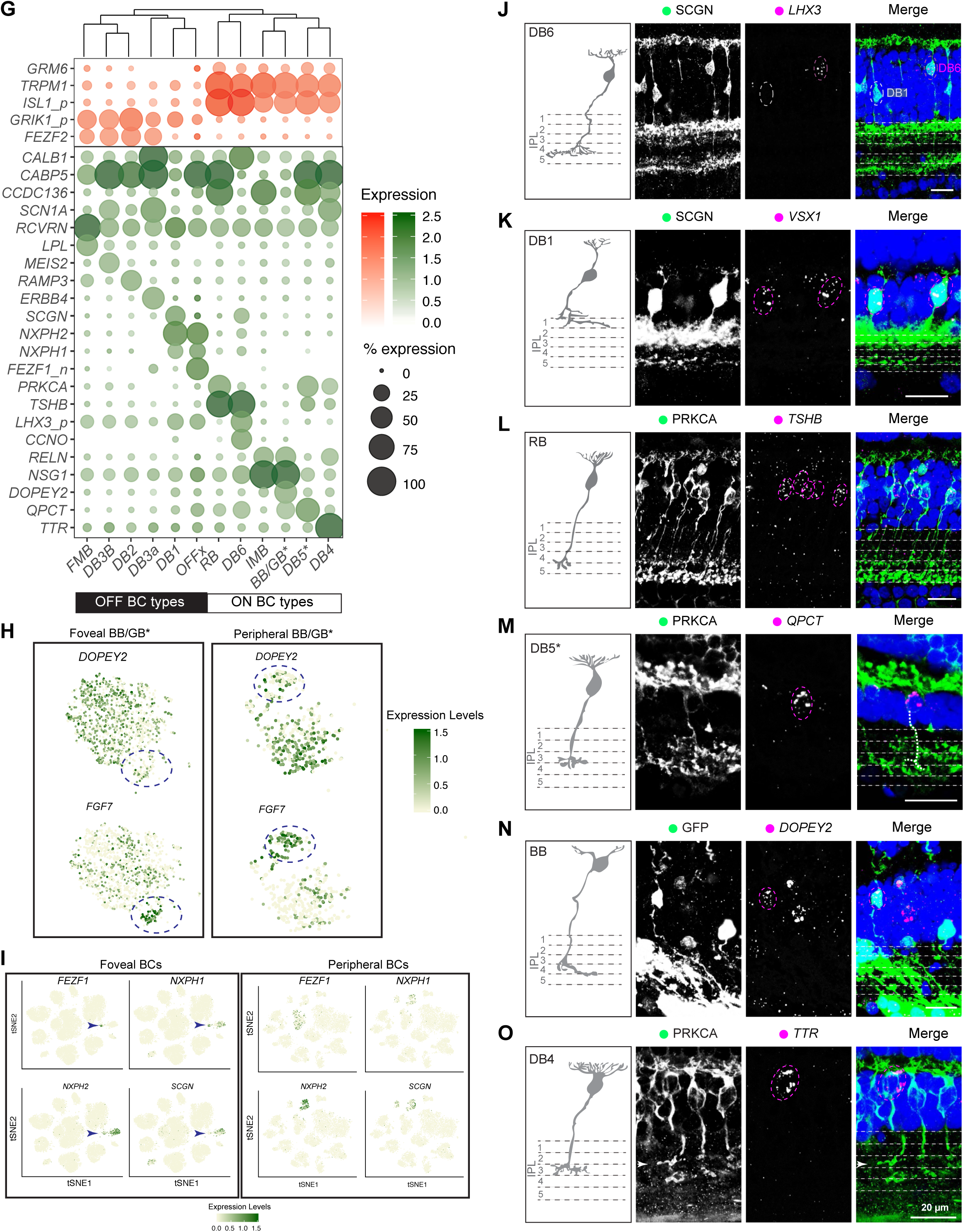
Characterization of cone and BC types. A. The reference macaque genome contains a single gene locus named *“OPN1MW/OPN1LW”* and, because of their high sequence similarity revealed from cloned cDNAs (98%; (Onishi et al., 2002)), our NCBI+TruSeq reference did not distinguish *OPN1MW* from *OPN1LW*. We therefore analyzed single-nucleotide polymorphisms (SNPs) to distinguish *OPN1MW-* and *OPN1LW*-specific expression in individual cones. An Integrated Genomics Viewer (IGV) screenshot of reads from a representative sample aligning to exon 3 of the loci shows that SNPs (arrows) are distinguishable from random sequencing errors at known locations (Onishi et al., 2002) based on their higher biallelic frequency. B. Counts of *OPN1LW*-specific SNPs vs *OPN1MW*-specific SNPs for every M, L cone (dots) show that most M/L cones in our data exclusively express either *OPN1MW* or *OPN1LW*. The remaining cells exhibit a mixed identity but only 0.8% show appreciable expression of both genes. It is generally believed that all M/L cones express either M or L opsin but not both. However, experimental evidence for this conclusion is lacking; the most authoritative studies, based on spectrophotometry of individual human cones in live retinae, cannot exclude the possibility that up to 5% of M/L express both opsins ((Hofer et al., 2005) and D. R. Williams, University of Rochester, personal communication). C. Proportions of M, L and mixed M/L cones in the fovea. D. Comparison of average transcriptional profiles of peripheral M-cones and L-cones identified through analysis of SNPs, as in **Figure 2A**. As in the fovea, no genes other than *OPN1MW* and *OPN1LW* differ significantly in expression levels (>1.2 fold at p<0.01, MAST test) between the two cone types. Dashed red line, y=x. As noted in the main text, the question of specificity in connections of M and L with synaptic partners has been controversial. If connectivity is selective, it is likely that M and L cones express different recognition molecules, implying transcriptional differences between the two that extend beyond the opsin genes. In addition to some physiological evidence for specificity (Reid and Shapley, 1992), there is a report of possible structural differences between the terminals of M and L cones (Lee et al., 2012). If wiring is nonselective no additional expression differences between the two cone types would be needed. Consistent with the idea, several physiological studies detect little (Field et al., 2010) or no selectivity (Wool et al., 2018) in chromatic inputs to MGC surrounds. A prevalent model is that the choice between expression of M and L opsin is stochastic, mediated by regulatory elements at the M/L locus (Wang et al., 1999). In this model no additional transcriptional differences would be predicted. Our results are consistent with this view. However, we cannot draw firm conclusions because it is possible that transcriptional differences are present during development but not maintained into adulthood. E. Proportions of M, L and mixed M/L cones in the peripheral retina. F. Additional FISH validations of markers that distinguish M/L-cones (*MYH4* and *VOPP1*) from S-cones (*TTR*) (see **Figure 2B**). Circles denote location of S cones in I. Scale bar is 20 μm. DAPI staining is in blue. G. Gene expression patterns (rows) of broad markers (top panel) and type-enriched markers (bottom panel) for foveal/peripheral BC types (columns) ordered using hierarchical clustering (see dendrogram on top). Representation as in **Figure 2D** H. *FGF7* and *DOPEY2* are enriched in distinct subpopulations within a single cluster that appears to contain both Blue Bipolar and Giant Bipolar types (BB/GB* cluster). Data were extracted from the full foveal and peripheral BC datasets and visualized by tSNE, as in **Figure 1F**. As *DOPEY2* (hi) cells correspond to the BB type (**Figure S2N**), we speculate that the *DOPEY2* (low), FGF7(hi) (dashed circles) subpopulation corresponds to the GB type. These assignments are consistent with their relative frequencies (**Table S2**). Colors represent expression levels. **Table S2** also includes references to prior studies that document type-specific expression of BC markers in primates. I. Expression of *FEZF1, NXPH1, NXPH2*, and *SCGN* in foveal (left) and peripheral BCs (right) visualized by tSNE as in **Figs.1F**. Coexpression of *FEZF1, NXPH1 and NXPH2* mark OFFx BCs in fovea (arrows), but they are not coexpressed in periphery, supporting the claim that OFFx may be a fovea-specific type. *SCGN* is expressed by DB1 and DB6 (see panels G, J, K) but not by OFFx. J. Secretogogin (SCGN) labels two BC types in the fovea, DB6 (magenta circle), which is also *LHX3+* (see **Figs. 2D,E**) and DB1 (white circle), which is *LHX3-*. K. DB1 (SCGN+) is also *VSX1+*. L., M. *TSHB* (L) and *QPCT* (M) label two distinct types of PRKCA+ BCs with distinct morphologies. Based on their abundance, *TSHB+* cells are rod bipolars (RB) and *QPCT+* cells could be DB5* (**Table S2**). The axon of DB5* was traced and is represented by a white dashed line. In contrast, PRKCA is a selective rod bipolar marker in mice (Shekhar et al., 2016) N. A viral labeled BB cell is *DOPEY2* positive. BB type has a distinguishable bifurcated dendritic morphology (Mariani, 1984). O. *TTR* is expressed by PRKCA+ DB4. Arrowheads indicate the DB4 axon terminal. In J-O, sketches were redrawn from (Tsukamoto and Omi, 2016). Scale bar, 20 μm. DAPI staining is in blue.

**Figure S3:**
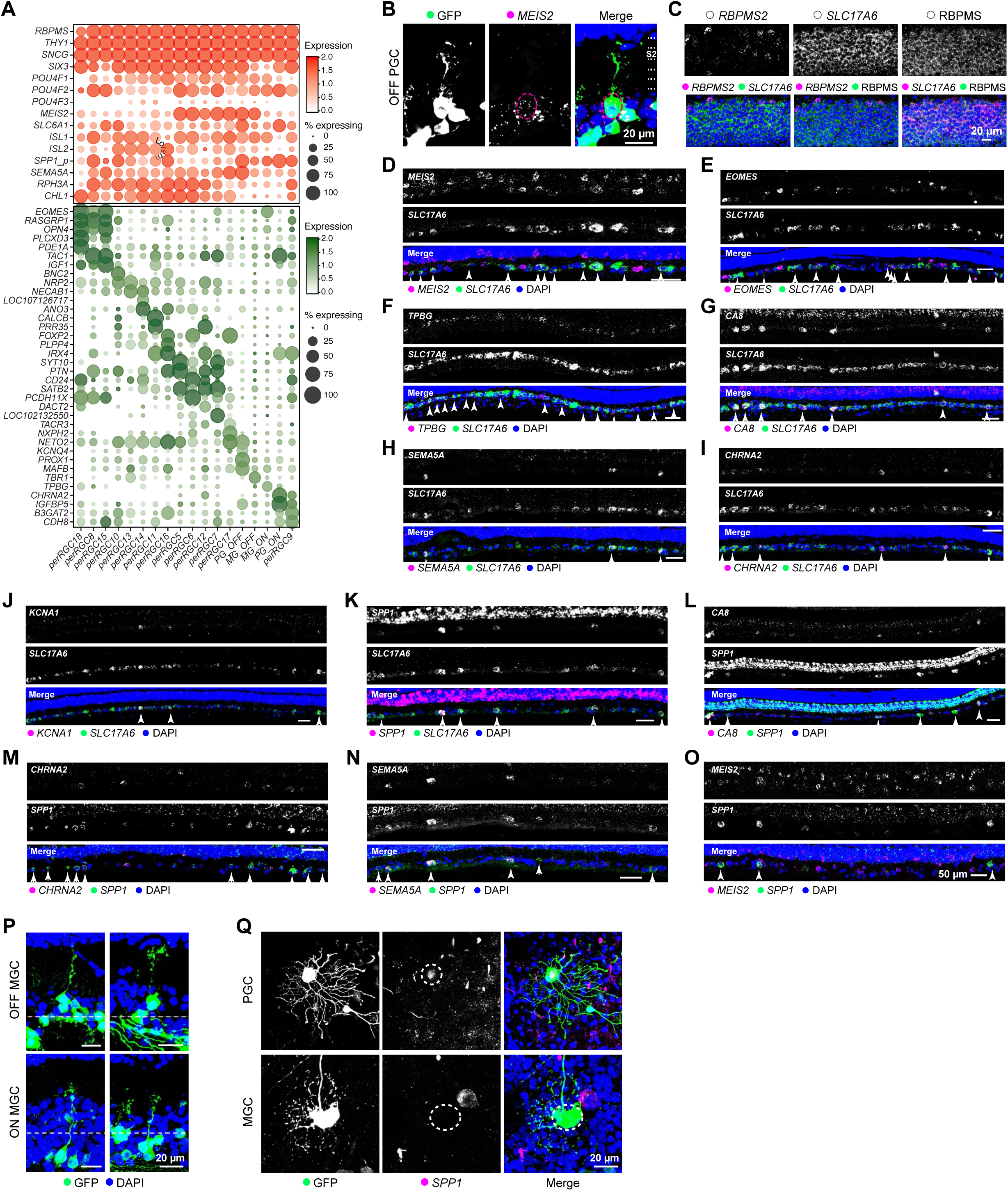
Histological validation of markers for RGCs. A. Gene expression patterns (rows) of broad (top panel) and type-enriched markers (bottom panel) for peripheral RGC clusters (columns) ordered by transcriptional similarity. Representation as in **Figure 2F**. See **Figure 3B** for transcriptional mapping of foveal and peripheral RGC clusters. B. FISH combined with viral labeling shows *MEIS2* expression by an OFF PGC with dendrites that arborize in the S1/S2 sublaminae of the IPL. C. Triple-labeling of a foveal section shows that all foveal RGCs are *SLC17A6+* and RBPMS+ (immunohistochemistry) but only a minority are *RBPMS2+*, consistent with its expression by PGCs but not MGCs (panel A). *SLC17A6* encodes VGlut2, which is a pan-RGC marker in rodents, but is not expressed by other cells in the ganglion cell layer (Stella et al., 2008). *RBPMS* is a selective marker for RGCs in both rodents and macaques (Rodriguez et al., 2014) but to our knowledge expression of its homologue, Rbpms2, has not been reported in mouse retina. D.-K., Double FISH shows that *MEIS2* (D), *EOMES* (E), *TPBG* (F), *CA8* (G), *SEMA5A* (H), *CHRNA2* (I), KCNA1 (J), and *SPP1* (K) colocalize with RGCs (*SLC17A6+*) in peripheral retina. Arrowheads indicate double positive cells. L.-O. Double FISH shows that *CA8* (L), *CHRNA2* (M), and *SEMA5A* (N) label subsets of *SPP1−* positive RGCs. In contrast, only a small fraction of *MEIS2+* RGCs are *SPP1+* (O); these are OFF PGCs (see panel B). Arrowheads indicate double positive cells; arrows indicate *SPP1* single positive cells. P. Viral labeling of foveal RGCs shows that the somata of OFF-MGCs (top) are localized at the outer half of the GCL while those of ON-MGCs are located at the inner half (bottom). Q. Viral labeling combined with immunostaining shows that a PGC cell expresses SPP1 (upper) but not a MGC (lower). Somata positions of labeled RGCs are circled in the middle panel. Cells are from the temporal peripheral retina. Scale bars is 20 μm (B, C, P, Q) and 50 μm (D-O).

**Figure S4:**
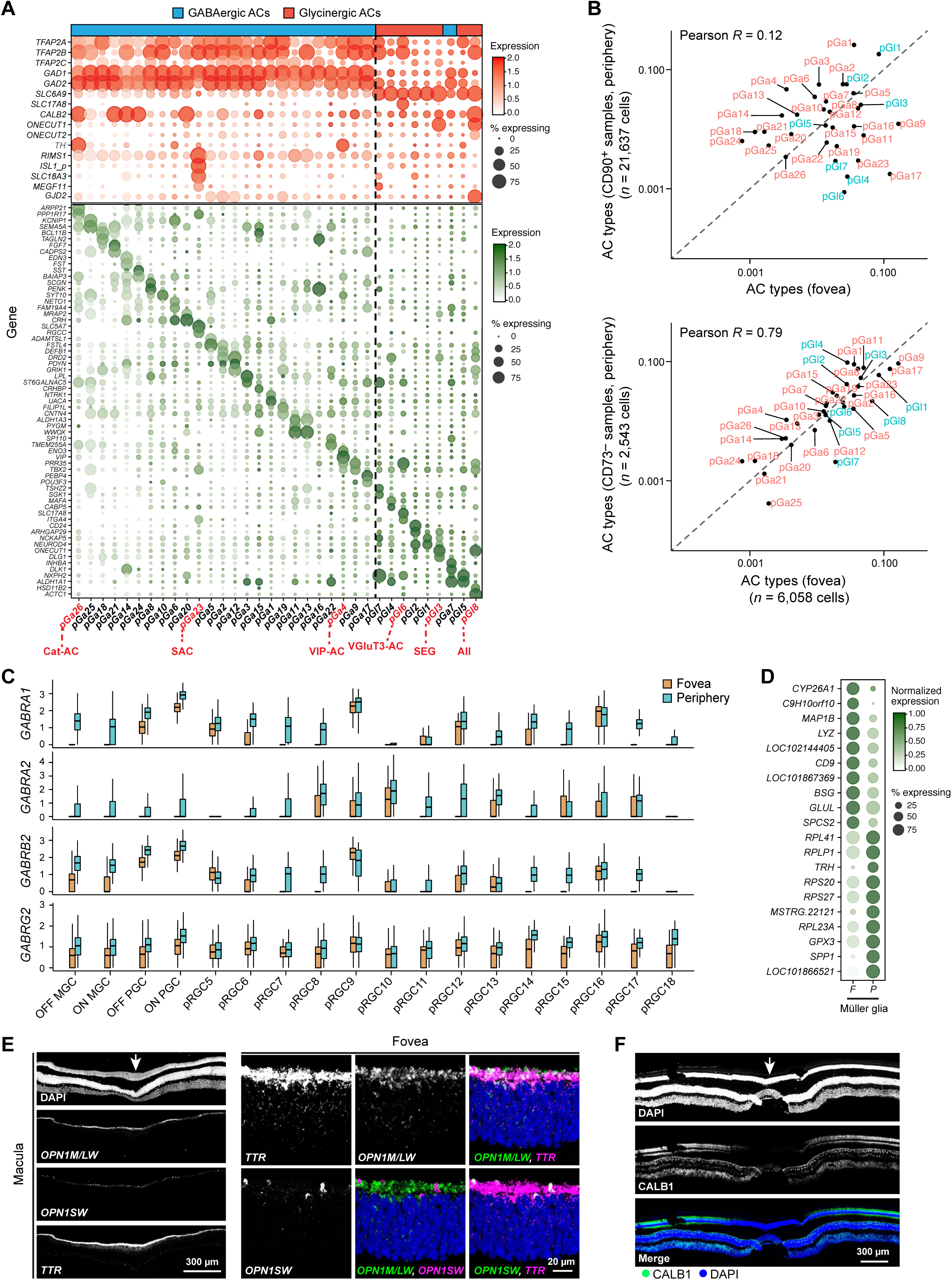
Differences in proportions of cell types and gene expression between the fovea and peripheral retina. A. Gene expression patterns (rows) of broad markers (top panel) and type-enriched markers (bottom panel) for 34 peripheral AC cluster (columns) ordered by transcriptional similarity, and their approximate segregation is highlighted by a black dashed line. Representation as in **Figure S2G**. Annotation bar on the top highlights GABAergic (n=26) and Glycinergic (n=8) types. Key known types of ACs including Starburst AC (SAC), AII ACs, Excitatory (VGluT3+) AC and Vasoactive Intestinal Peptide expressing (VIP+) AC, catecholaminergic (Hendrikson et al.) and SEG AC are highlighted below. Rodent AC types have been described in detail (Diamond, 2017; Haverkamp and Wassle, 2004; Kay et al., 2011; Krishnaswamy et al., 2015; Park et al., 2015; Zhang et al., 2007) as have several primate equivalents (Klump et al., 2009; Lammerding-Koppel et al., 1991; Majumdar et al., 2008; Mariani and Hokoc, 1988; Mills and Massey, 1999; Yamada et al., 2003). B. Comparison of frequencies of matched AC types between the foveal (x-axis) and peripheral datasets (y-axis). To facilitate comparison, we assigned each foveal AC a peripheral identity based on **Figure 3C** and then compare frequencies of 1:1 matched types. The peripheral ACs are subdivided into those originating from CD90+ samples (upper) and CD73− samples (lower). Values are plotted on a ln-ln scale to highlight low frequency types. The poor correlation of the AC types in the CD90+ sample with the fovea suggests a biased labeling of AC types by this marker. Hence, we only used AC cluster frequencies in the CD73− samples, which show a much higher correlation to the foveal frequencies, to compute compositional similarities between the fovea and the periphery. C. Expanded version of **Figure 4F**, showing higher expression levels of GABR_A_ receptor subunits *GABRA1/2, GABRB2* and *GABRG2* in the majority peripheral RGC compared to foveal RGC types. For 1:1 comparison each foveal RGC is assigned a peripheral identity based on **Figure 3B**. D. Gene expression differences between foveal and peripheral MGs. The average number of transcripts per expressing cells (colors) are normalized so as to sum up to 1 along each row to highlight relative differences. E. Co-labeling of *TTR* with both *OPN1LW* and *OPN1SW* in the fovea suggests that it is expressed by both foveal M/L and S cones. F. Foveal cones were CALB1-negative, as documented in (Hendrickson et al., 2007). Scale bars, 20 μm (E), and 300 μm (E,F). Arrows indicate the center of fovea In the bar and whisker plots in C, black horizontal line, median; bars, interquartile range; vertical lines, minimum and maximum. Scale bars is 20 μm. DAPI staining is blue in A and B.

**Figure S5:**
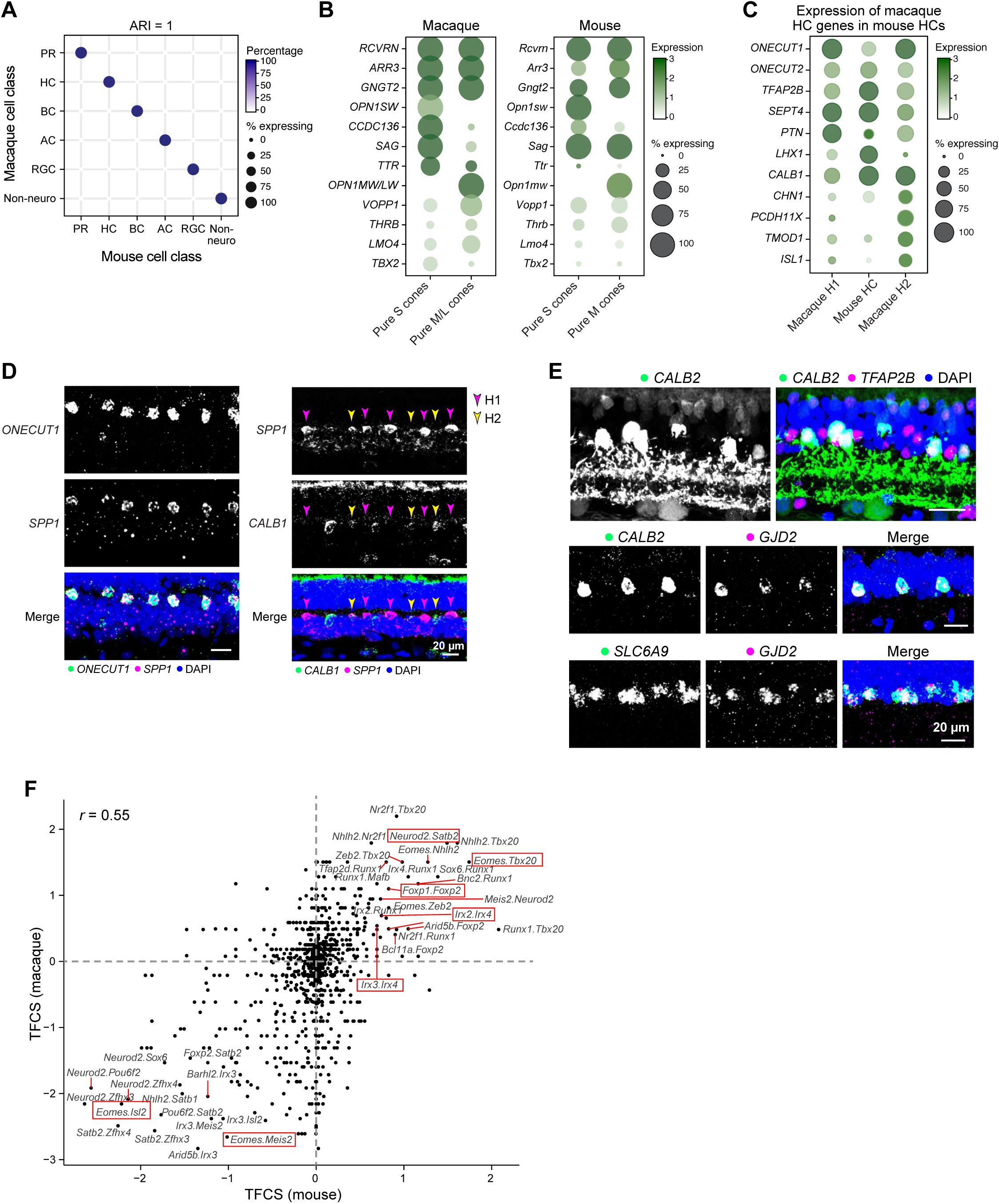
Comparison of mouse and macaque cell types. A. Transcriptional correspondence between macaque (rows) and mouse (columns) cell classes. All types are included within each class. Representation as in **Figure 5A** B. Mouse S cones express higher levels of *Ccdc136* compared to M cones, the mouse ortholog of macaque S cone marker *CCDC136* (**Figure 2B**). Other S cone or M/L cone markers in macaque, however, are not shared in mice. *CCDC136* locus is located near the *OPN1SW* locus in both mouse and human (~16 kb separation on human chromosome 7 and mouse chromosome 6) suggesting the possibility of transcriptional co-regulation, possibly by NRL (Brooks et al., 2011). *THRB*, selectively expressed by OPN1MW-expressing mouse cones and M/L macaque cones, has been implicated in differentiation of this cone type (Ng et al., 2001). C. Mouse HCs are transcriptionally more similar to macaque H1 (**Figure 5B**), and do not express markers that are specific to macaque H2. D. SPP1 is a macaque HC marker gene and H2 and H2 are distinguished by the expression of CALB1. Double FISH of *SPP1* and *CALB1* in a peripheral retina section labels both H1 (*SPP1+CALB1-*, magenta arrows) and H2 (*SPP1+CALB1+*, orange arrows). E. Validations of conserved AII markers: (Top) AII ACs, identifiable by their unique morphology, are CALB2+TFAP2B+. Double FISH shows that coexpression of AII marker *GJD2* (Macosko et al., 2015) with *CALB2* (middle) and *SLC6A9* (bottom). F. Comparison of transcription factor co-expression scores (TFCS, see **Methods**) of pairs of transcription factors for macaque (y-axis and mouse (x-axis) RGC types. Dashed red lines denote the x and the y axes. The relative predominance of points in the (+,+) and (−,−) quadrants compared to the (+,−) and (−,+) quadrants suggests that synergistic and antagonistic relationships between TFs are largely preserved across the species. Red boxes highlight several examples in which TF pairs show similar patterns of co-expression or mutually exclusive expression in both species. Expression of transcription factors in mouse RGC subsets is documented in (Cherry et al., 2011; Liu et al., 2018; Mao et al., 2014; Peng et al., 2017; Rousso et al., 2016; Sweeney et al., 2017). Scale bar, 20 μm

**Figure S6:**
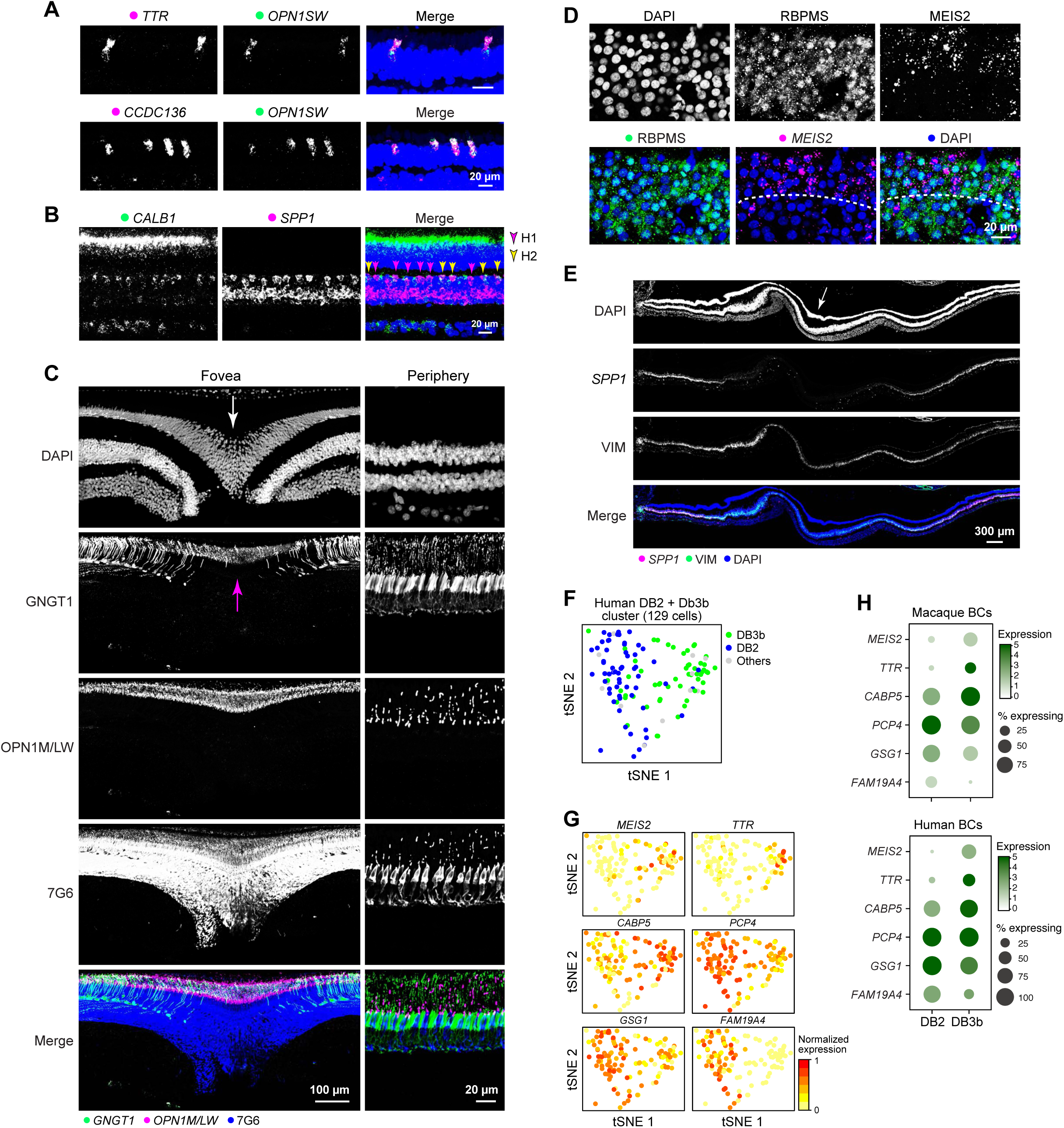
Key molecular features identified from the macaque retinal atlas are conserved in marmosets and humans. A. *TTR* and *CCDC136* positive S cones in marmoset B. H1 (*SPP1+CALB1−*) and H2 (*SPP1+CALB1+*) HC in the marmoset peripheral retina. C. Foveal cones are GNGT1 positive in the marmoset. Staining marmoset foveal and peripheral retinal tissues with antibodies against GNGT1, OPN1LW, and 7G6 (cone arrestin) shows that GNGT1 is specifically expressed by foveal cones and localized at the outersegments (indicated by the expression of OPN1LW) of foveal cones. Arrows indicate the center of fovea. D. The expression of *MEIS2* (outer) in the human foveal GCL layer. E. *SPP1* (F) is expressed by peripheral Müller glia (*VIM*) respectively in human retina. Arrows indicate the center of fovea. F. tSNE plot of cells from human bipolar cluster DB2+3b, color coded by the macaque peripheral bipolar type that each human cell mapped to. Gray indicating cells mapped to clusters other than DB2and DB3b. G. Expression of DB2 and DB3b genes in the DB2+3b cluster. H. Expression of the same gene sets as G in macaque and human peripheral bipolar cells Scale bar, 20 μm (A, B, C, D), 100 μm (C), and 300 μm (E). DAPI staining is in blue.

**Figure S7:**
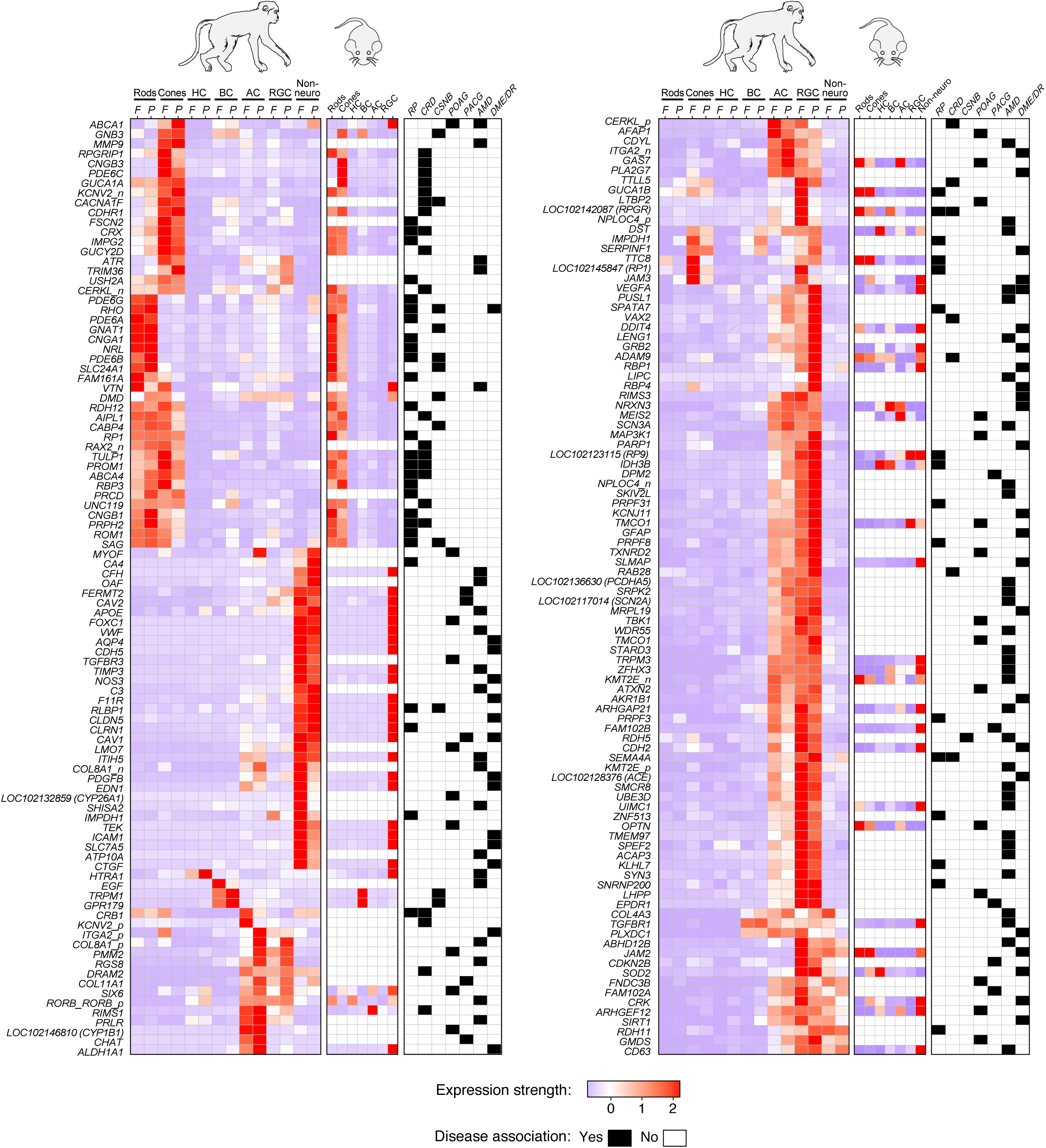
Expression Patterns of Retinal-Disease Associated Genes across Major Cell Classes in macaque fovea and periphery, as well as mouse retina. Red-blue heatmaps show expression patterns of individual retinal-disease associated genes (rows) by cell classes (columns), for macaque and mouse respectively, as **Figure 7B**. For each gene, associated retinal diseases determined by GWAS studies are shown in white-black heatmaps. For genes in the macaque transcriptome that have been mapped, but not characterized (e.g *LOC102123115*), their predicted human orthologs are indicated in parenthesis.

## Footnotes

1 https://www.ncbi.nlm.nih.gov/genome/776

